# Temperature stress resilience in polar *Chlamydomonas* is regulated by acclimation to light and salinity: implications for survival in a changing world

**DOI:** 10.64898/2026.04.03.716389

**Authors:** Pomona Osmers, Allison Szenasi, Laura Kostyniuk, Sophia Caputo, Nicholas Bradette, Marina Cvetkovska

## Abstract

- Aquatic algae are key primary producers in the Arctic and Antarctic, yet how cold-water species respond to environmental change is poorly understood. The Polar Regions are increasingly exposed to frequent heat waves, leading to declining ice cover, increased light availability, and decreasing salinity in polar waters. We compared three phylogenetically related but geographically distant polar *Chlamydomonas* species to test how habitat history shapes algal responses to light, salinity, and temperature stress.
- We assessed the growth, morphology, and photochemistry of psychrophilic *Chlamydomonas* acclimated to native-like (lower light, higher salinity) and climate-shifted conditions (higher light, lower salinity). Next, we exposed acclimated cultures to a lethal heat shock and observed how acclimation affects algal temperature stress resilience.
- All three species acclimated to climate-shifted conditions grew rapidly but showed the greatest sensitivity to temperature stress, with rapid loss of viability and photosynthetic efficiency. In contrast, slow-growing cultures acclimated to native-like conditions exhibited significantly greater resilience to temperature stress.
- Our work is the first to directly link light and salinity acclimation with temperature resilience in psychrophilic algae, suggesting that fast-growing polar green algae may be particularly vulnerable to increasingly frequent heat waves, with major implications for primary productivity in polar environments.

## Introduction

The Arctic and Antarctic are the largest ecosystems on our planet, but the global cryosphere is shrinking. Surface air temperatures in the polar regions have increased 2-4x more than the global average, resulting in the loss of >28 trillion tonnes of ice since 1994 (Slater *et al*., 2021). Modeling studies predict an alarming possibility of an ice-free Arctic Ocean within the 21^st^ century (Stroeve & Notz, 2018; Wunderling *et al*., 2020) and there are reports of increased frequency and length of heatwaves, even in the interior of the Antarctic continent (e.g., 18.3°C; Turner *et al*., 2021; Feron *et al*., 2021). Primary production in the Polar Regions is largely dependent on photosynthetic microbes, including eukaryotic algae from all major lineages (Anderson & Macdonald, 2015; Bax *et al*., 2021). Many photosynthetic algae endemic to the Polar Regions are psychrophiles that thrive in the cold (0-5°C) but can not withstand even very modest temperature increases (Morita, 1975; Cvetkovska *et al*., 2017). Temperature sensitivity and loss of ice habitats make psychrophiles particularly threatened by current climate trends.

Light and salinity are key drivers in polar ecosystem dynamics and are particularly important for the growth of photosynthetic algae. Thinning ice and increased duration of seasonally open waters allow for an increase in light availability that drives photosynthesis in aquatic habitats (Obryk *et al*., 2019; Castellani *et al*., 2022). Greater solar penetration within the water column also contributes to warming and increased melting at the ice-water interface below the surface, resulting in runaway ice-cover disintegration (Bers *et al*., 2013; Cárdenas *et al*., 2018; Lehnherr *et al*., 2018; Sherwell *et al*., 2022). The resulting rapid influx of glacial meltwater leads to global “freshening” of marine waters, an effect that is particularly pronounced in the Polar Regions (Durack & Wijffels, 2010; Silvano *et al*., 2018; Bronselaer *et al*., 2018). These environmental changes are accompanied by well documented ecosystem-scale increases of primary production and greening of both poles (Lehnherr *et al*., 2018; Myers-Smith *et al*., 2020a; Williamson *et al*., 2020; Gray *et al*., 2020; Dalpadado *et al*., 2020; Roland *et al*., 2024). Despite this apparent positive effect on algal growth, little is known about the species composition of these microbial assemblages and the fate of endemic psychrophiles within them.

The order Chlamydomonadales (Chlorophyta) harbours more than a third of all confirmed algal psychrophiles, with a wide distribution in Antarctic, Arctic, and alpine habitats (Cvetkovska *et al*., 2017). This group includes several well-described green algae with sequenced genomes that have been elevated to the status of models for photosynthetic life at the extreme (Cvetkovska *et al*., 2019; Zhang *et al*., 2019, 2020, 2021). Among these, the Antarctic *Chlamydomonas priscui* (formerly UWO241; Morgan-Kiss & Guiry, 2024) is the only psychrophile that has been studied in the context of temperature stress. Exposure to acute non-permissive temperature (24°C) results in alterations in global metabolomic and transcriptomic responses after 6h (Cvetkovska *et al*., 2022b), followed by a rapid drop in photosynthetic rates after 12h, and cellular death with prolonged exposure (Possmayer *et al*., 2011). These studies suggest that psychrophilic algae have a limited ability to induce a heat stress response (HSR) similar to that of their mesophilic relatives (Schroda *et al*., 2015), insufficient in overcoming their inherent temperature sensitivity.

These breakthrough studies opened the door to understanding psychrophilic responses to environmental stress, but *C. priscui* is unique even among polar Chlamydomonadales. This alga is endemic to the deep photic zone of the Antarctic Lake Bonney in the McMurdo Dry Valleys (Neale & Priscu, 1995), where the perennial 3-6 m thick ice cover severely restricts external inputs, resulting in a highly stable and stratified environment characterized by strict temperature, light, salinity, and nutrient profiles (Patriarche *et al*., 2021). In its native environment, *C. priscui* experiences permanently low temperatures (∼4°C), very low light (∼10 μmol m^-2^ s^-1^) biased in blue-green wavelengths (400-500 nm), high salinity (∼700 nm NaCl), high O_2_ saturation (30-40 mg L^-1^) and low nutrient levels (Neale & Priscu, 1995). It has been postulated that life at such stable conditions has resulted in a ‘locked’ physiology that confers adaptive advantages in an extreme and stable habitat, but lacks the capacity to respond to short term environmental challenges (recently reviewed in Cvetkovska *et al*., 2022a; Hüner *et al*., 2022, 2024). Therefore, while *C. priscui* is undoubtedly the best characterized polar green alga, its responses to environmental stress may not be representative of psychrophiles in general.

Psychrophilic *Chlamydomonas* have been detected in virtually all polar habitats (Cvetkovska *et al*., 2017), but our knowledge on their stress resilience capabilities are largely unknown. Open marine waters, sea ice, and snowfields are populated by a wide diversity of photosynthetic microbes (Balzano *et al*., 2012; van Leeuwe *et al*., 2018; Davey *et al*., 2019; Gérikas Ribeiro *et al*., 2020; Dorrell *et al*., 2023) that experience annual freeze-thaw cycles accompanied by dramatic light, salinity, and temperature fluctuations (Tremblay *et al*., 2012; Urbański & Litwicka, 2022; Spolaor *et al*., 2024). This natural seasonal variability might influence the ability of psychrophilic algae to acclimate to new conditions, and we hypothesized that polar algae originating from a dynamic environment will have an increased ability to withstand environmental stress.

To address this, we compared the physiology of three phylogenetically related but geographically distant psychrophilic *Chlamydomonas* species in the face of acute and sustained environmental challenge. In addition to *C. priscui*, we focused on two of its relatives: 1) the euryhaline *Chlamydomonas malina* native to the open waters of the Arctic Beaufort Sea that experience seasonal freeze-thaw cycles, deeper light penetration, and frequent vertical mixing (Taylor *et al*., 2013); and 2) the freshwater *Chlamydomonas klinobasis* isolated from ephemeral snowfields on Spitsbergen Island in the Arctic (Hulatt *et al*., 2017), a dynamic oligotrophic environment characterized by high light intensities on the surface and rapid light attenuation within the snow pack (Stibal *et al*., 2007). Using light and salinity as variables, we characterized the physiological acclimation strategies of these species to climate change scenarios and their responses to subsequent temperature stress. This work provides a unique opportunity to compare the stress resilience of closely related species that originate from cold but otherwise very different environments and enhances our understanding of climate-sensitive psychrophilic algal biology.

## Materials and Methods

### Algae cultivation and experiment design

*Chlamydomonas priscui* (formerly UWO241; CCMP1619, https://ncma.bigelow.org/) was isolated 17 meters below the ice surface in Lake Bonney, Antarctica (Neale & Priscu, 1995).

*Chlamydomonas malina* (RCC2488; https://roscoff-culture-collection.org/) was isolated from the surface waters of the Beauford Sea in the Canadian Arctic (Balzano *et al*., 2012). *Chlamydomonas klinobasis* (CCCRyo 050-99; https://cccryo.fraunhofer.de) originates from a snowfield on Spitsbergen in the Svalbard archipelago (Hulatt *et al*., 2017).

Axenic algal stocks were maintained on Bold’s Basal Media (BBM) agar slants at 4°C with continuous illumination (∼50 μmol m^-2^ s^-1^). Working liquid stocks were seeded from agar-grown cultures in 250 mL Erlenmeyer flasks in BBM media. Experimental cultures were grown in custom-made closed culture bioreactors suspended in temperature-regulated aquaria and aerated with sterile ambient air provided by aquarium pumps. Experimental cultures were acclimated to two different steady-state light intensities and salinities. Continuous light (10 µmol m^-2^ s^-1^ or 100 µmol m^-2^ s^-1^) was provided by full-spectrum LED bulbs. Intensity was modulated with neutral density filters and monitored with a quantum light sensor (Model LI-189; Li-COR). Salinity was modified by supplementing BBM media with NaCl at the appropriate concentration (10 mM and 700 mM for *C. priscui*; 10-600 mM for *C. malina*; 0.43-70 mM for *C. klinobasis*). All cultures were allowed to acclimate to these conditions for at least 4 weeks before experimentation. For temperature stress experiments, algae were cultured at 4°C until the mid-exponential growth phase (OD_750_ = 0.3-0.4). Stress was initiated by moving the cultures to a non-permissive temperature: 24°C for *C. priscui* and 22°C for *C. klinobasis* and *C. malina*. All experiments were done on cultures during mid-exponential growth and with at least three biological replicates (independently seeded cultures), each with four technical replicates (individual bioreactor tubes).

### Phylogenetic analysis

To determine the phylogenetic relationship between the three species, we constructed gene trees using the nuclear-encoded 18S *rRNA* and plastid-encoded *RbcL* genes. The sequenced genomes of *C. priscui* (Zhang *et al*., 2021) and *C. klinobasis* (JGI Project Id: 1106927; https://genome.jgi.doe.gov/portal/ChlkliStandDraft_3) were used to source the *RbcL* and 18S *rRNA* nucleotide sequences. To obtain RbcL sequences from *C. malina*, tissue was collected during the mid-exponential phase and total DNA was isolated as previously described (Zhang *et al*., 2021). Gene fragments were amplified using universal green algae *RbcL* primers (Forward: 5’-GGTACTTGGACAACAGTTTGGAC -3’; Reverse: 5’-TACGCAACCTCTCTGGCAAAG-3) (Hadi *et al*., 2016), and the amplicon was sequence using Sanger sequencing (Génome Québec, Montreal, Canada). All other sequences were obtained from GenBank with a BLASTn search within the Chlamydomonadales (taxid: 3042).

To ensure appropriate representation, we used the sequences from the species studied here as queries, as well as sequences from model algae. The data were manually curated to remove duplicate and low-quality sequences. To avoid overrepresentation, we removed sequences with ≥98% identity that originated from the same species but different strains. Using this approach, we obtained all available sequences from the Monadinia and Moewusinia clades (sensu Nakada; (Nakada *et al*., 2008) and from selected representatives from other Chlamydomonadales clades. This resulted in a total of 64 *RbcL* and 111 18S *rRNA* sequences used in downstream analyses. All sequences were aligned using MAFFT (Katoh & Standley, 2013) and trimmed to remove gaps and ambiguously aligned regions with BMGE (Criscuolo & Gribaldo, 2010). Phylogenetic relationships were inferred using maximum likelihood in PhyML v.3.0 (Guindon *et al*., 2010) via the NGPhylogeny platform (Lemoine *et al*., 2019). The best-fitting model was selected using the Smart Model Selection (SMS) based on the Akaike Information Criterion (AIC) (Lefort *et al*., 2017). Tree topology searches were performed using the SPR algorithm, and statistical support for branches was evaluated using 500 non-parametric bootstrap replicates. Trees were rooted at species belonging to the Sphaeropleales, a sister clade to the Chlamydomonadales (Buchheim *et al*., 2001; Lemieux *et al*., 2015), and annotated in iTOL (Letunic & Bork, 2021).

### Growth and viability measurements

Algal growth was monitored by measuring pigment content, cell counts, and optical density (OD). To measure pigment content, 1 mL of cultures was pelleted by centrifugation (3 min, 16,000 rcf), the supernatant was removed and replaced with 90% (v/v) acetone. The pellet was re-suspended and centrifuged again to remove cellular debris (3 min, 16,000 rcf). The absorbance of the supernatant was measured spectroscopically at 467, 664 and 647 nm (Cary 60, Agilent Technologies). Chlorophyll (Chl) *a* and *b* content was calculated according to Jeffrey & Humphrey (1975). Carotenoid (Car) content was calculated according to Sun *et al*. (1998), with the absorbance maximum shifted from ß-carotene to astaxanthin. Cell numbers were quantified with an automatic cell counter (Countess II FL, ThermoFisher Scientific) using brightfield imaging (EVOS^TM^ Light Cube, white) and Chl autofluorescence imaging (EVOS^TM^ Cy5 Light Cube, Ex/Em = 628/685 nm). Change in OD was measured at 750 nm (Cary 60, Agilent Technologies). Growth rates were calculated using OD_750_ values during exponential growth.

Cell viability was quantified by labelling algal cells with the fluorescent dye SYTOX Green (Ex/Em = 504/523 nm; ThermoFisher Scientific) that accumulates in dead cells only. SYTOX Green dissolved in DMSO was added to algal samples at a final concentration of 2 μM, followed by 15 min dark incubation prior to fluorescent imaging. Green fluorescence, corresponding to dead cells, was measured using an automatic cell counter (EVOS™ GFP Light Cube, Ex/Em=470/525 nm). Cell death was quantified as a proportion of green fluorescing cells to the total number of cells. To quantify the loss of Chl (bleaching) during stress, pigments were extracted as above, and bleaching was calculated as a percentage of the Chl content immediately before stress initiation.

### Flow cytometry and microscopy

Cellular size and complexity was examined using a flow cytometer (Gallios, Beckman Coulter). In each case, SYTOX Blue (Ex/Em = 405/480 nm; ThermoFisher Scientific) dissolved in DMSO was added to algal culture (∼1×10^6^ cells/mL) at a final concentration of 1 µM, and incubated in the dark for 5 minutes with gentle agitation at the experimental temperature. Data were collected as at least 20,000 events for each sample, at a flow rate of 500 events s^-1^. Algal cells were identified through Chl autofluorescence using the FL7 detector (725 ± 20 nm), thus eliminating cellular debris or contaminants from downstream analyses. Dead cells stained with SYTOX Blue were detected by the FL9 detector (450 ± 40 nm) and excluded from further analyses.

Data was analyzed using the Kaluza Analysis Software (Beckman Coulter, v2.1). Cell size and granularity were obtained from the forward scatter (FS) and side scatter (SSC) measurements, respectively. The ratio between FS height and FS area was used to determine cell size and estimate the proportion of single cells and palmelloids. These measurements were further confirmed by measuring FS and SSC to estimate internal complexity, expected to be higher in multi-cell palmelloid colonies. The cellular morphology was further confirmed by the presence of two distinct chlorophyll autofluorescence peaks exhibiting minimal overlap, as demonstrated previously (Szyszka-Mroz *et al*., 2022). Algae were visualized using brightfield imaging on a Zeiss AxioImager microscope using the Zen Pro Blue V. 3.2.0 software (Carl Zeiss AG).

### Room temperature chlorophyll a fluorescence

*In vivo* room temperature Chl *a* fluorescence was measured using a pulse amplitude modulated chlorophyll fluorometer (DUAL-PAM-100; Walz) with the WinControl software. Live cultures (1 mL; 2 μg Chl/mL) were stirred with a magnetic stir bar and kept at constant temperature (4°C or 22-24°C) using a water-jacketed quartz cuvette connected to a circulating water bath. After 10 minutes of dark adaptation, Chl fluorescence at open PSII reaction centers (F_o_) was excited by a non-actinic measuring light (0.34 μmol m^-2^ s^-1^) pulsed at 20 Hz. Maximum fluorescence at closed PSII reaction centers in the dark (F_m_) was induced by a saturating pulse (200 ms, 10,286 μmol m^-2^ s^-1^). To measure maximal fluorescence at closed PSII reactions centers (F_m_’) and the steady-state fluorescence in the light (F) saturating flashes were applied in 30 s intervals under constant actinic light of 27 μmol m^-2^ s^-1^. All photosynthetic parameters were calculated according to (Kramer *et al*., 2004). Maximum PSII photochemical efficiency (F_V_/F_M_) was calculated as F_M_–F_o_/F_M_, using dark-adapted cells. The quantum yield of steady-state photosynthesis in the light was calculated as: Y(II) = (F_M_′–F_S_)/F_M_′, Y(NPQ) = (F/F_M_′)-(F/F_M_), Y(NO) = F/F_M_, where Y(II) is the yield of PSII photochemistry, Y(NPQ) is the yield of non-photochemical energy dissipation by down-regulation through antenna quenching, and Y(NO) is the yield of all other processes involved in non-photochemical energy losses. The relative electron transport rate (ETR) at PSII was determined as ETR = Y(II) x (PDF x 0.84) x 0.5.

### Statistical analysis

All statistical analyses were conducted in R using the rstatix, car, and emmeans packages (www.r-project.org). Growth rates, pigment quantification, and cell sizes were analyzed by two-way ANOVA and Tukey’s post-hoc test, with salinity and light intensity as independent variables. Cellular viability was analyzed using linear models and Tukey’s post-hoc test with salinity, light intensity, and time as independent variables. Photosynthetic measurements were analyzed by three-way repeated measures ANOVA with Dunett’s post-hoc test, using salinity and light intensity as independent variables.

## Results

### The phylogeny and morphology of psychrophilic Chlamydomonas

The phylogenetic analysis of the *RbcL* and 18S *rRNA* sequences positioned all three psychrophilic *Chlamydomonas* species used in this work within the Monadinia and Moewusinia clades of the Clamydomonadales. The topologies of the phylogenetic trees are consistent with previously published data (Nakada *et al*., 2008, 2016; Lemieux *et al*., 2015). The phylogenetic position of the Antarctic *C. priscui* in a unique lineage within the Moewusinia clade was previously described (Possmayer *et al*., 2016). Here, we report that the closest known relative of this alga is the Arctic *C. malina*, with a high bootstrap support (99%, Fig. 1, Fig. S1, S2) and sharing a close phylogenetic relationship with *Chlamydomonas* sp. WAP05 and WAP30 isolated from the Southern Ocean (Soto *et al*., 2023). Our analysis provides strong support (87-96%) for the placement of the snow alga *C. klinobasis* in the same clade as *Microglena monadina* (Demchenko *et al*., 2012) and other species from the polar regions (e.g., *Chlamydomonas* sp. ICE-L. ICE-W, ICE-MDV, KNF0022) (Liu *et al*., 2006; Eddie *et al*., 2008; Jung *et al*., 2016; Cook *et al*., 2019) (Fig. 1, Fig. S1, S2).

**Figure 1:**
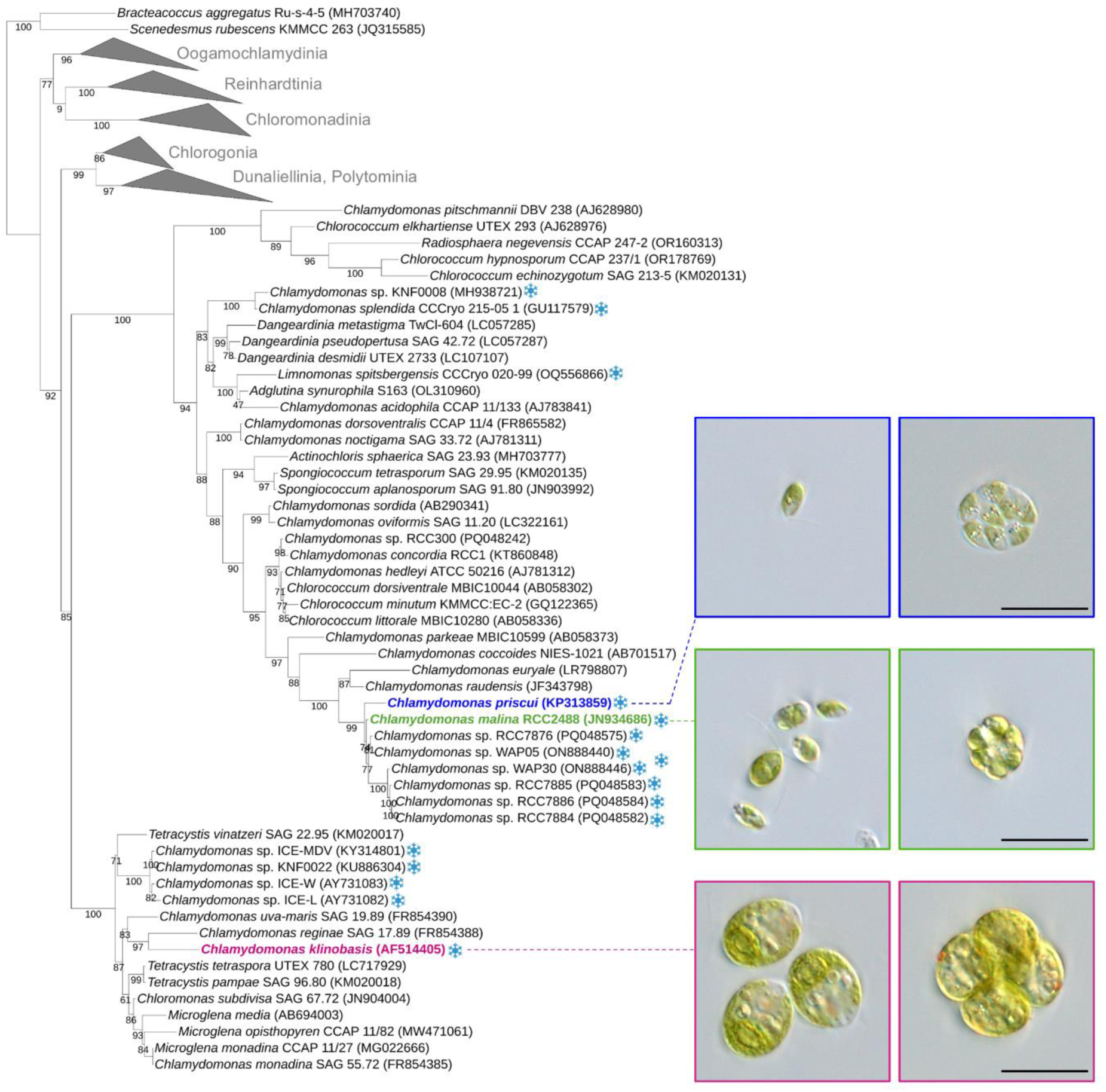
Phylogenetic tree of 111 18S rRNA nucleotide sequences inferred from using maximum likelihood analysis is shown to scale (substitutions per site), with branch values representing bootstrap support. GenBank accession numbers are provided in brackets beside the organism name. Collapsed clades contain select green algal species within major *Chlamydomonas* taxa and are represented by triangles at the branch names, while the Monadinia and Moewusinia clades are shown in full. Algae isolated from polar environments are indicated with a blue symbol. The species considered in this study are highlighted in blue (*Chlamydomonas priscui*), green (*Chlamydomonas malina*) and mauve (*Chlamydomonas klinobasis*). The images in the inset show light micrographs of these species as single cells (left) and palmelloid colonies (right). Scale bar = 10 µm.

Morphologically, all three species exist as biflagellate free-swimming single cells on non-motile palmelloid colonies, where multiple single cells are encased in an outer limiting membrane (Fig. 1, inset). The occurrence of these morphological variants has been described in *C. priscui* (Pocock *et al*., 2004; Possmayer *et al*., 2016; Szyszka-Mroz *et al*., 2022), and here we showed that single cells and palmelloids also co-occur in *C. malina* and C*. klinobasis* cultures. As previously reported (Pocock *et al*., 2004; Possmayer *et al*., 2016), *C. priscui* single cells were ellipsoidal or sub-fusiform, approximately 6-7 μm long, with a single cup-shaped chloroplast, and very small eyespot. The morphology of *C. malina* was very similar with ovoid, ellipsoidal or sub-fusiform cells with a single chloroplast, albeit smaller in size (∼4-6 μm long) and displaying a more prominent eyespot compared to *C. priscui* (Fig. 1, inset). Finally, single cells in the freshwater *C. klinobasis* were subspherical or ovoid and significantly larger (∼12-13 μm in length) compared to the halotolerant species, with a very prominent eyespot visible in both single cells and palmelloids (Fig. 1, inset).

### Salinity and light affect the growth and physiology of psychrophilic Chlamydomonas

The three psychrophilic *Chlamydomonas* species used here originate from geographically and environmentally distinct habitats. Thus, we monitored the growth and morphology of these algae at environmentally relevant (rather than identical) conditions (Table 1). For the halotolerant *C. priscui* and *C. malina*, the high salinity (HS) treatment was chosen to mimic their natural habitat (700 mM and 300 mM, respectively) (Smoot & Hopcroft, 2017; Patriarche *et al*., 2021). For the freshwater *C. klinobasis*, the HS treatment (30 mM) was determined experimentally to represent the salinity that supports robust growth comparable to that of *C. priscui* and *C. malina* at high salinity (Fig. S3a; Table S1). Conversely, the low salinity treatment for *C. klinobasis* (LS, 0.43 mM) was chosen as an approximation of salinities measured in snow (Tonboe *et al*., 2021; Saha *et al*., 2025). For the halotolerant species, low salinity was defined as the salinity at which these algae exhibited the highest growth rates and comparable to those of *C. klinobasis*, previously reported as 10 mM for *C. priscui* (Possmayer *et al*., 2011; Cvetkovska *et al*., 2022b) and experimentally determined as 30 mM for *C. malina* (Fig. S3b; Table S1). For all three species (Table 1), the low light treatment (LL, 10 µmol m^-2^ s^-1^) represents the intensity received by polar algae in their natural habitat under ice or a snow cover (Neale & Priscu, 1995; Stibal *et al*., 2007; Hill *et al*., 2018). The high light treatment (HL, 100 µmol m^-2^ s^-1^) mimics light availability in shallow open waters (Hill *et al*., 2018; Sherwell *et al*., 2022) and is the recommended intensity for culturing various *Chlamydomonas* species in the lab (Hui *et al*., 2023). All three species had comparable growth rates when acclimated to these experimental conditions (Table S1).

**Table 1:**
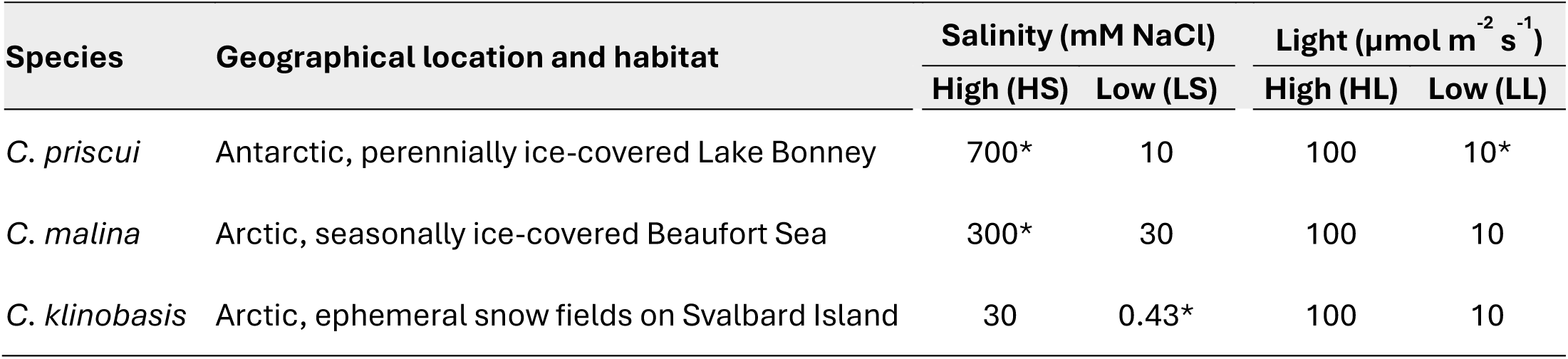
Algal species, their geographical location, and the experimental conditions used in this work. The salinity and light intensity that resemble each species natural habitat are emphasized with (*). While C. priscui originates from a stable environment characterized by low light availability, *C. malina* and *C. klinobasis* originate from dynamic environments and experience variable light conditions.

Light availability was a major driver of growth in all three species, but only in *C. malina* and *C. klinobasis* cultures growth was also dependent on salinity or the interaction between the two variables (Fig. 2a, Table S2). *C. priscui* cultures achieved maximal growth when acclimated to higher light intensity (0.35-0.49 ΔOD_750_ day^-1^) compared to those acclimated to low light (0.19-0.26 ΔOD_750_ day^-1^), regardless of salinity (Fig. 2a). In contrast, *C. malina* and *C. klinobasis* cultures exhibited highest growth rates with higher light availability and lower salinity (0.71 and 0.47 ΔOD_750_ day^-1^, respectively), and decreased growth rates when cultured at higher salinities (0.45 and 0.28 ΔOD_750_ day^-1^, respectively). Both species grew significantly slower when cultured at lower light intensity, regardless of the salinity (0.21-0.23 ΔOD_750_ day^-1^ for *C. malina*; 0.15-0.23 ΔOD_750_ day^-1^ for *C. klinobasis*) (Fig. 2a).

**Figure 2:**
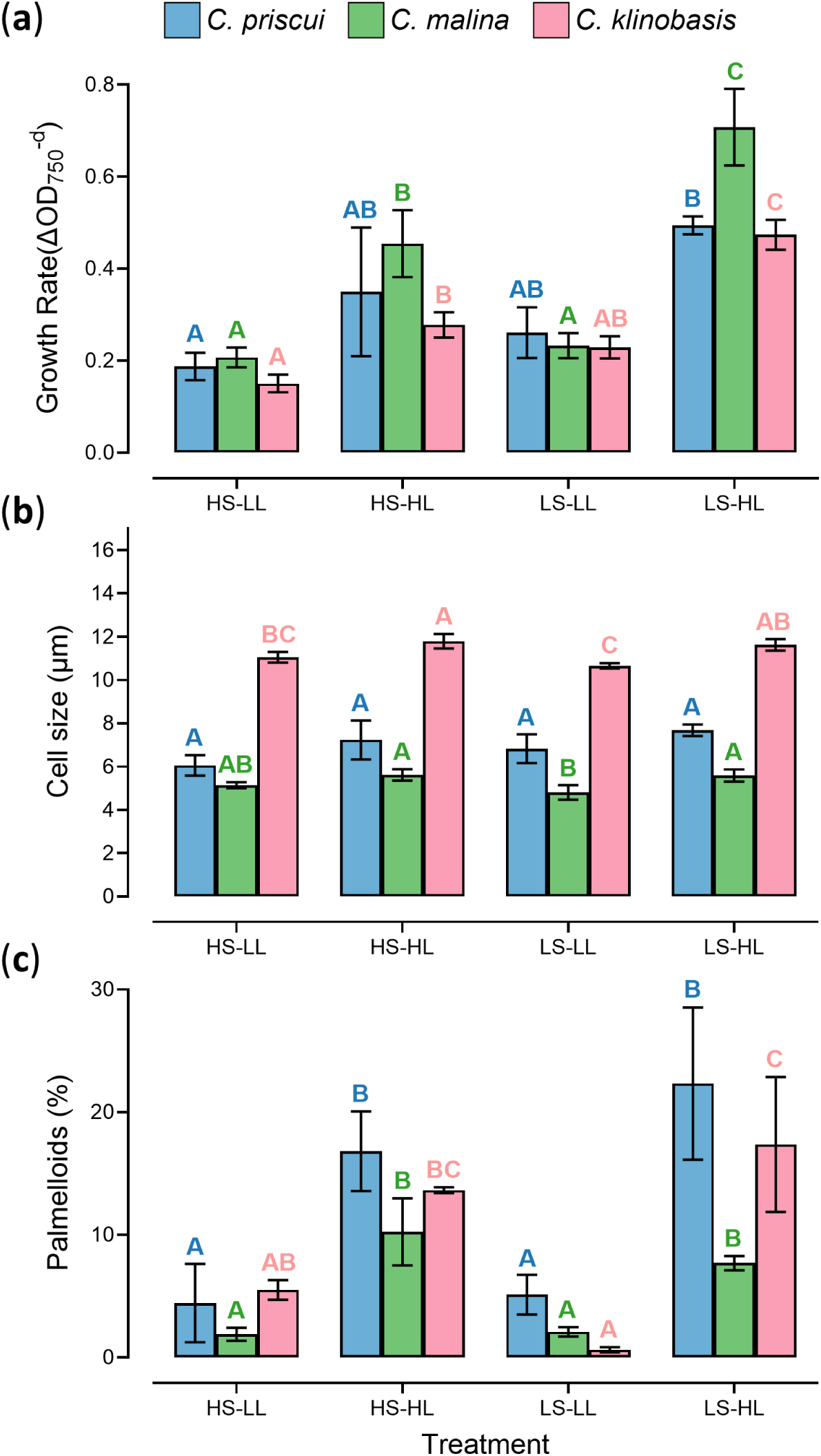
The effects of salinity and light on the growth and morphology of *C. priscuii*, *C. malina* and *C. klinobasis*. (**a**) Maximal growth rates. (**b**) Average cell size. (**c**) A percentage of cells present as palmelloid colonies. In all cases, algae were cultured at 4°C and acclimated to high salinity (HS), low salinity (LS), high light (HL) or low light (LL). Data are means ± SD of at least three biological replicates. Statistically significant differences in the parameters measured within each species were determined with two-way ANOVA with a Tukey’s post-hoc test (p<0.05) and are represented as different letters.

All three species can exist as single motile cells or multi-cell palmelloids (Fig. 1), and we observed that light availability (but not salinity) has a significant effect on algal morphology (Table S2). While cell size (∼6.9 μm) did not change significantly in the lacustrine *C. priscui* cultures in response to experimental conditions (Fig. 2b), the palmelloid content was ∼4x higher at cultures acclimated to high light compared to those at lower light intensities (Fig. 2c). The marine *C. malina* had significantly larger single cells (∼5.6 μm) and ∼4x more palmelloid colonies when cultured with higher light availability, in comparison with cultures grown at lower light (∼5.0 μm) (Fig, 2b, 2c). Finally, the snow alga *C. klinobasis* had the largest cells, with significant cell size difference and 5x higher palmelloid content between high light (∼11.7 μm) and low light (∼10.9 μm) acclimated cultures (Fig. 2b, 2c).

Light availability was also a major environmental contributor of steady-state pigment content in all three species, but we demonstrated that salinity has a species-specific effect (Table 2; Table S2). All three species accumulated ∼1.5-2x more Chl/cell when acclimated to low light intensity compared to cultures grown with higher light, regardless of salinity (Table 2). While *C. klinobasis* typically accumulated 2-4x more chlorophyll compared to *C. malina* and *C. priscui*, this is likely a reflection of the larger cell size in this species (Fig. 2b). Acclimation to limited light led to lower Chl *a/b* ratios in all species, but salinity had a significant effect only in *C. malina* cultures. This Arctic alga grown at higher salinity exhibited small but significant (∼7%) increase in Chl *a/b* ratios compared to cultures grown at lower salinity, when acclimated to equivalent light levels (Table 2, Table S2). Car/Chl ratios were also regulated by light in all three species, where algae acclimated to higher light exhibited a higher ratio of the two pigments (Table 2). While salinity had no effect on the pigment composition balance in *C. malina*, Car/Chl ratios were affected by both variables in *C. priscui* and *C. klinobasis*. In both cases, cultures acclimated to high salinity accumulated more carotenoids compared to those acclimated to low salinity at the equivalent light, leading to a significant decrease (5-13%) in the Car/Chl ratios (Table 2).

**Table 2:** The effects of salinity and light on the pigment content of *C. priscui*, *C. malina* and *C. klinobasis*. In all cases, algae were cultured at 4°C and acclimated to high salinity (HS), low salinity (LS), high light (HL), or low light (LL). Data are means ± SD of at least three biological replicates. Statistically significant differences within each species were determined with two-way ANOVA with a Tukey’s post-hoc test (*p* < 0.05) and are represented as different letters.

|  | HS-LL | HS-HL | LS-LL | LS-HL |
| --- | --- | --- | --- | --- |
| ***C. priscui*** |  |  |  |  |
| **Chl/cell (pg)** | 3.10±0.58<sup>a</sup> | 2.43±0.57<sup>a</sup> | 3.66±0.96<sup>a</sup> | 2.30±0.2<sup>a</sup> |
| **Chl a/b** | 2.03±0.09<sup>b</sup> | 3.08±0.15<sup>a</sup> | 2.18±0.07<sup>b</sup> | 3.28±0.21<sup>a</sup> |
| **Car/Chl** | 0.41±0.01<sup>b</sup> | 0.44±0.01<sup>a</sup> | 0.39±0.01<sup>c</sup> | 0.42±0.01<sup>ab</sup> |
| ***C. malina*** |  |  |  |  |
| **Chl/cell (pg)** | 2.21±0.25<sup>a</sup> | 1.15±0.12<sup>b</sup> | 2.17±0.31<sup>a</sup> | 1.07±0.05<sup>b</sup> |
| **Chl a/b** | 1.92±0.03<sup>d</sup> | 3.02±0.07<sup>b</sup> | 2.06±0.02<sup>c</sup> | 3.26±0.07<sup>a</sup> |
| **Car/Chl** | 0.45±0.01<sup>b</sup> | 0.51±0.01<sup>a</sup> | 0.44±0.01<sup>b</sup> | 0.5±0.01<sup>a</sup> |
| ***C. klinobasis*** |  |  |  |  |
| **Chl/cell (pg)** | 5.01±0.29<sup>ab</sup> | 3.62±0.57<sup>b</sup> | 9.51±3.79<sup>a</sup> | 4.44±0.83<sup>a</sup> |
| **Chl a/b** | 2.67±0.08<sup>b</sup> | 4.18±0.05<sup>a</sup> | 2.61±0.05<sup>b</sup> | 3.92±0.37<sup>a</sup> |
| **Car/Chl** | 0.46±0.01<sup>c</sup> | 0.6±0.02<sup>a</sup> | 0.42±0.01<sup>d</sup> | 0.52±0.01<sup>b</sup> |

Chl, chlorophyll; Car, carotenoids

### Rapid growth correlates with increased sensitivity to temperature stress

Culturing at different salinity and light levels affects growth, morphology, and pigment content in *polar Chlamydomonas* (Fig. 2, Table 2) but how acclimation to these conditions affects algal responses to temperature stress has not been examined before. To test this, we exposed algae acclimated to four different conditions (Table 1) to long-term temperature stress. To compare the temperature sensitivity of the three species, we used an equivalent stress model instead of an equal temperature model. *C. priscui* is the only psychrophilic green algae that has been examined under temperature stress (24°C) (Possmayer *et al*., 2011; Cvetkovska *et al*., 2022b). Thus, we tested the sensitivity of *C. malina* and *C. klinobasis* to a range of non-permissive temperatures and selected those where the kinetics of chlorophyll bleaching and cell death most closely mirrored those previously established in *C. priscui*. We show that both *C. malina* (Fig. S3c) and *C. klinobasis* (Fig. S3d) cultures incubated at 22°C lose viability at similar rates as *C. priscui* cultured at 24°C, when acclimated to equivalent light and salinity levels. This suggests that these two species, despite being isolated from dynamic environments, are more sensitive to temperature stress compared to *C. priscui*.

We demonstrate that acclimation to different environmental conditions has a significant effect on the resilience of psychrophilic algae to long-term temperature stress (Fig. 3). In all three species, temperature-induced cell death (Fig. 3a, 3b, 3c) and chlorophyll loss (Fig. 3d, 3e, 3f) occurred most rapidly in cultures acclimated to low salinity and higher light intensity (3-4 days to 100% viability loss; 4-5 days to 100% chlorophyll loss). These are the conditions that led to the fastest growth rates prior to exposure to temperature stress (Fig. 2a). Conversely, slow-growing cultures acclimated to high salinity and low light intensity (Fig. 2a) exhibited robust resilience to temperature stress and gradually lost viability (Fig. 3a-c) and chlorophyll (Fig. 3d-f) over ≥15 days at increased temperature.

**Figure 3:**
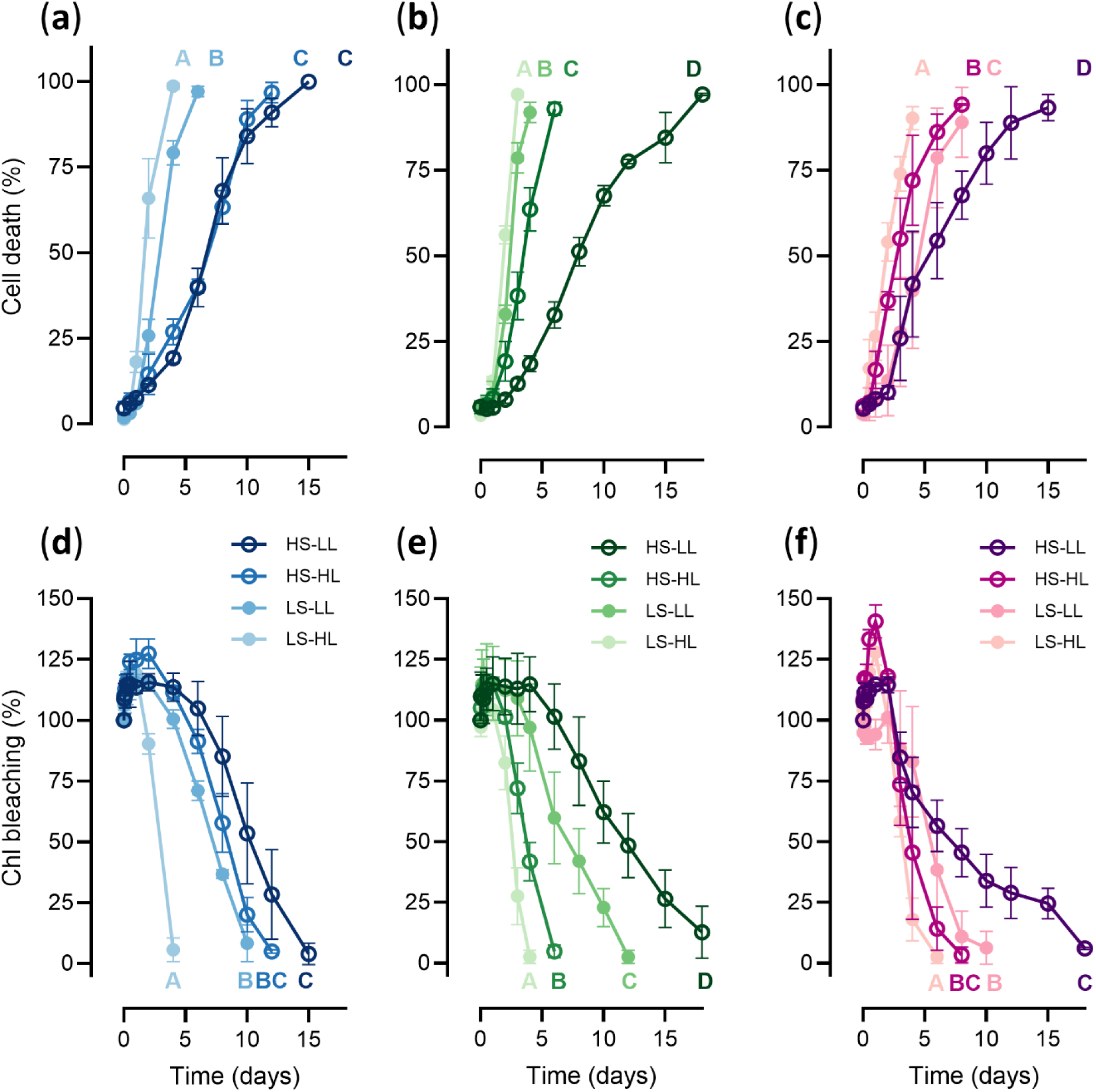
The kinetics of cell death (**a, b, c**) and chlorophyll bleaching (**d, e, f**) in *C. priscui* (**a, d**; blue), *C. malina* (**b, e**; green) and *C. klinobasis* (**c, f**; mauve) exposed to non-permissive temperature. In all cases, algae were cultured at 4°C and acclimated to high salinity (HS), low salinity (LS), high light (HL) or low light (LL). Temperature stress was initiated at the mid-exponential growth phase (OD_750_ = 0.3-0.4) and corresponds to t = 0 days. Cell death was calculated as a percent of algal cells stained with the live-dead stain SYTOX Green. Chlorophyll bleaching was calculated as the proportion of chlorophyll present at each time point compared to levels prior to exposure to temperature stress (0 days). Data are means ± SD of at least three biological replicates. Statistically significant differences in the parameters measured within each species were determined with two-way ANOVA with a Tukey’s post-hoc test (*p*<0.05) and are represented as different letters.

In addition to these robust trends, the interaction between light and salinity had a more subtle, species-specific effect. Salinity was the main driver behind temperature resilience in the lacustrine *C. priscui*, where cultures acclimated to 700 mM NaCl exhibited slow cell death and chlorophyll loss kinetics, regardless of the light intensity (Fig. 3a, 3d; Table S3). Light intensity had a significant effect on the kinetics of chlorophyll loss in this alga, but only in cultures acclimated to low salinity (Fig. 3d; Table S3). Both salinity and light intensity significantly affected the viability of the marine *C. malina* exposed to temperature stress. Cultures acclimated to low salinity prior to temperature stress suffered more rapid heat-induced cell death compared to those acclimated to higher salinity. Conversely, higher light-acclimated cultures lost their chlorophyll content faster during heat stress, compared to those acclimated to low light (Fig. 3e; Table S3). Finally, light intensity appeared to be the main factor behind temperature sensitivity in the snow alga *C. klinobasis*. Algae acclimated to higher light exhibited more rapid rates of cell death and chlorophyll loss compared to those grown at lower light, regardless of the salinity (Fig. 3c; Fig. 3f; Table S3).

### Acclimation to light and salinity affects PSII photochemistry during temperature stress

Photosynthesis is highly sensitive to environmental stress and is affected before chlorophyll loss or cell death become apparent (Nievola *et al*., 2017; Lodeyro *et al*., 2021). All three psychrophilic species had robust photosynthetic capacity in all conditions when grown at 4°C. In all cases, F_V_/F_M_ (0.7-0.8; Fig. 4a-c) and ETR (11-14 μmol m^-2^ s^-1^; Fig. 4d-f) values were within the normal range for green algae (Malapascua *et al*., 2014). The exception was *C. priscui* acclimated to high salinity, with lower F_V_/F_M_ (0.52-0.56; Fig. 4a) and ETR (8-9 μmol m^-2^ s^-1^; Fig. 4d). We also observed lower F_V_/F_M_ values (0.64-0.67; Fig. 4c) in *C. klinobasis* cultures acclimated to higher light intensity, although this alga maintained high ETR rates at these conditions (11.2-13.3 μmol m^-2^ s^-1^; Fig. 4f).

**Figure 4:**
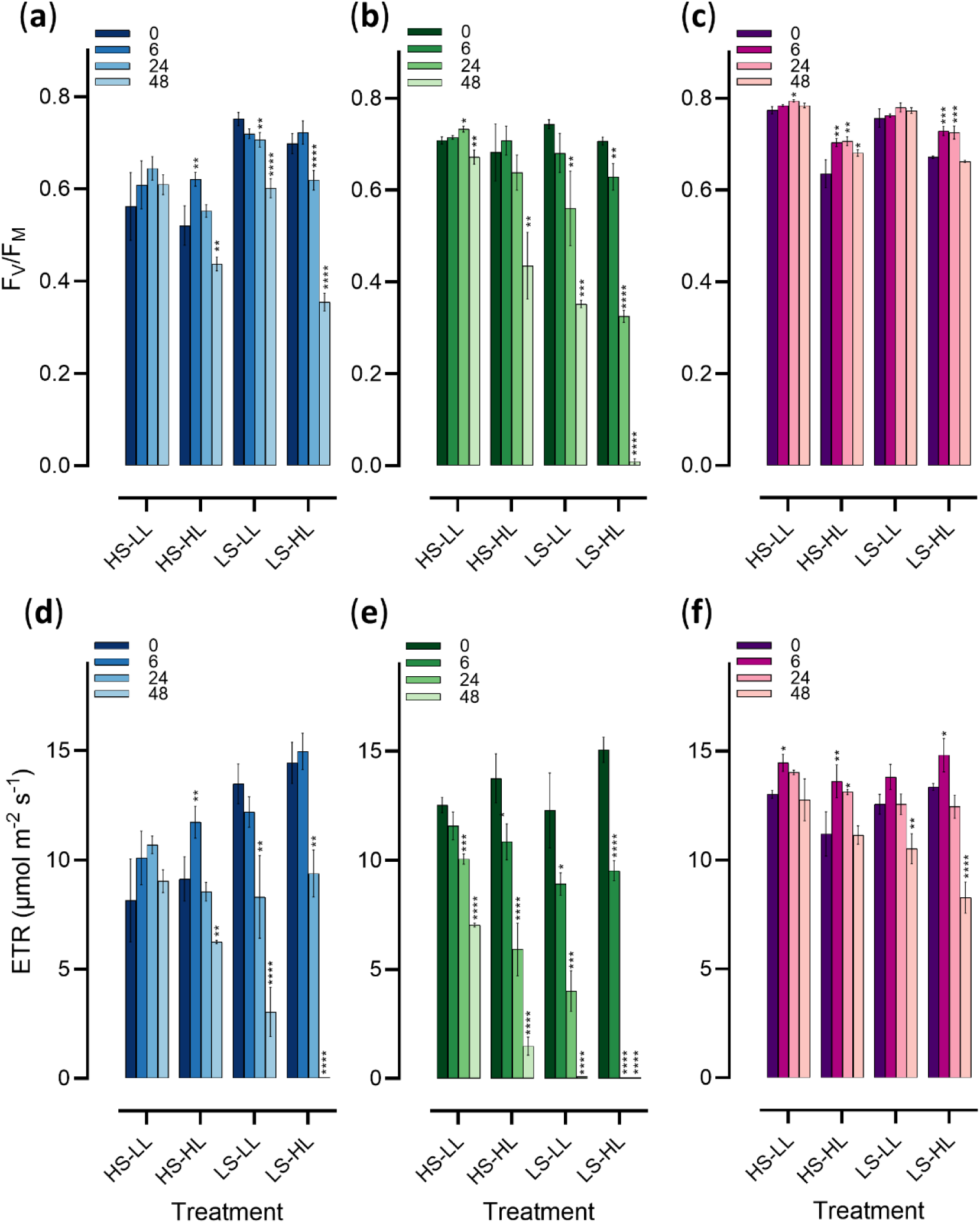
The maximum PSII photochemical efficiency (F_V_/F_M_; **a, b, c**) and relative electron transport rate (ETR; **d, e, f**) in *C. priscui* (**a, d**; blue), *C. malina* (**b, e**; green) and *C. klinobasis* (**c, f**; mauve) exposed to non-permissive temperature. In all cases, algae were cultured at 4°C and acclimated to high salinity (HS), low salinity (LS), high light (HL) or low light (LL). Temperature stress was initiated at the mid-exponential growth phase (OD_750_ = 0.3-0.4) and corresponds to t = 0 hours. Data are means ± SD of at least three biological replicates. Statisti cal significance compared with the value prior to stress application (t=0) within each species were determined with three-way ANOVA with a Dunnett’s post-hoc test (* = < 0.05, ** = < 0.01, *** = < 0.001, **** = < 0.0001). Detailed statistical analyses can be found in Table S3

Salinity and light intensity significantly affected photochemistry during temperature stress in all three species (Table S3), but the magnitude of this effect depended on the initial acclimation conditions. Mirroring chlorophyll loss and cell death trends, photosynthetic parameters measured in slow-growing algal cultures acclimated to high salinity and low light were the least affected by temperature stress. *C. priscui* (Fig. 4a, 4d) and C*. klinobasis* (Fig. 4c, 4f) cultures did not exhibit s decrease in F_V_/F_M_ and ETR over 48 hours at increased temperature. Instead, we observed a small increase in these parameters during the first 24 hours, although these values are significant only in *C. klinobasis* (35% increase in F_V_/F_M_ at 24h; 10% increase in ETR at 6h). *C. malina* was the most sensitive to temperature increase at the level of the photosynthetic apparatus with significant decreases in both parameters (5% decrease in F_V_/F_M_, Fig. 4b; 44% decrease in ETR, Fig. 4e) after 48 hours of temperature stress.

Fast growing cultures acclimated to low salinity and higher light were the most sensitive to temperature stress. Both *C. priscui* (Fig. 4a) and *C. malina* (Fig. 4b) experienced a significant decrease in maximum quantum efficiency after 48 hours (49% and 99% decrease in ΔF_V_/F_M_, respectively). Rates of electron transport were severely affected and not detectable after 48 hours in *C. priscui* (Fig. 4b) and only 24 hours in *C. malina* (Fig. 4e). These species acclimated to a combination of high salinity and high light, or low salinity and low light, exhibited similar decreases in F_V_/F_M_ and ETR, albeit to a smaller extent (Fig. 4a, 4b, 4d, 4e). *C. klinobasis* was the least affected by temperature stress. While ETR rates were moderately decreased by 48 hours of temperature stress in cultures acclimated to low salinity and higher light (32%, Fig. 4f), F_V_/F_M_ exhibited a transient increase during this period (7%; Fig. 4c).

Light energy absorbed by PSII can be used for photochemistry [Y(II)] or dissipated, either through regulated processes and antenna quenching [Y(NPQ)] or through non-regulated processes [Y(NO)] (Kramer *et al*., 2004). Our results indicate that energy partitioning is sensitive to temperature increases in polar *Chlamydomonas*, and highly dependent on the salinity and light acclimation prior to stress (Fig. 5; Table S3). Incubation at non-permissive temperature caused a significant increase in Y(NPQ) and Y(NO) at the expense of Y(II) in both *C. priscui* and *C. malina* at 48h, a response that was dependent on the acclimation to both light and salinity (Table S3). The exception to this trend occurred in *C. priscui* acclimated to high salinity and low light, where Y(II) increased after 48 hours of stress (10%; Fig. 5a; Table S4). Acclimation to all other growth conditions led to decreases in Y(II), although the effect was moderate (32% decrease) in cultures grown at high salinity and high light, and severe in those acclimated to low salinity (77-100% decrease; Fig. 5a, Table S4). The Arctic *C. malina* suffered significant stress-induced decrease in Y(II) (Fig. 5b, Table S4). While this decrease was moderate in algae acclimated to high salinity and low light (44%), we observed a precipitous decline in all other conditions after 48 hours (∼89-100%). The decline was particularly severe in algae grown at low salinity and high light intensity, with a complete loss of photochemical quenching by 24 hours of temperature stress (Fig. 5b, Table S4).

**Figure 5:**
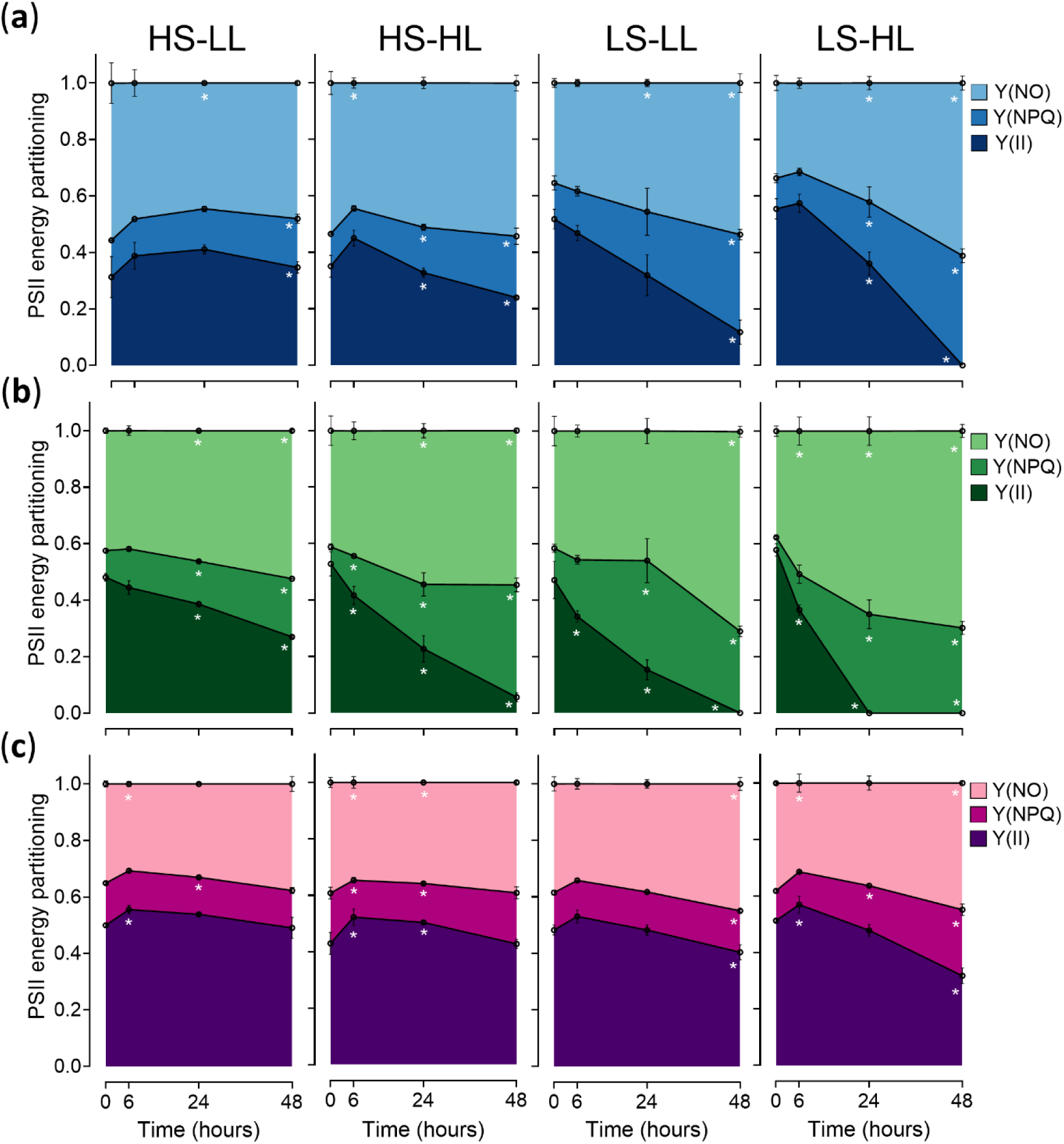
Energy partitioning at PSII in *C. priscui* (**a**; blue), *C. malina* (**b**; green), and *C. klinobasis* (**c**; mauve) exposed to non-permissive temperature. The total absorbed light is distributed between photochemical yield of PSII Y(II), non-photochemical quenching Y(NPQ), and non-regulated dissipation Y(NO). In all cases, algae were cultured at 4°C and acclimated to high salinity (**HS**), low salinity (**LS**), high light (**HL**) or low light (**LL**). Temperature stress was initiated at the mid-exponential growth phase (OD_750_= 0.3-0.4) and corresponds to t = 0 hours. Data are means ± SD of at least three biological replicates. Statistically significance compared with the value prior to stress application (t=0) within each species were determined with three-way ANOVA with a Dunnett’s post-hoc test (* = < 0.05). Detailed statistical analyses are available in Table S3 and S4.

Finally, photochemistry in the snow alga *C. klinobasis* was the least affected by temperature stress, compared to its halotolerant relatives. Incubation at non-permissive temperature caused a rapid but transient increase in Y(II) by 6 hours (10-18%) accompanied by a decrease in Y(NO) (12-13%) in all culturing conditions (Fig. 5c, Table S4). Photochemical quenching was negatively affected by 48 hours only in the cultures acclimated to low salinity (16% and 38% at low and high light respectively; Fig. 5c, Table S4).

## Discussion

### Close relatives from opposite sides of the globe

Our phylogenetic analysis places all three *Chlamydomonas* species examined here in the Moewusinia and Monadinia clades within the Chlamydomonadales, an order that represents one of the most taxonomically and phylogenetically complex groups of green algae (Nakada *et al*., 2008; Demchenko *et al*., 2012). The taxonomic identity of the model Antarctic psychrophile *C. priscui* has experienced much change (Morgan-Kiss and Guiry, 2024) but thorough phylogenetic analysis placed this alga in unique lineage within the Moewusinia clade (Possmayer *et al*., 2016). A close phylogenetic relationship with the marine *Chlamydomonas parkeae* (SAG 24.89) suggested a mesophilic ancestor prior to the arrival of *C. priscui* in Lake Bonney (Possmayer *et al*., 2016). Here, we demonstrate that the Arctic marine *C. malina* (RCC2488, Balzano *et al*., 2012; Morales-Sánchez *et al*., 2020) and the Antarctic marine C. sp. WAP05/30 (Soto *et al*., 2023) are the closest known relatives of *C. priscui* (Fig. 1, Fig. S1, S2), raising other possibilities for the ancestry of this group. The Arctic snow alga *C. klinobasis* also shares a close relationship with both Arctic and Antarctic psychrophiles within the Monadinia clade (Fig. 1, Fig. S1, S2), including *Chlamydomonas* sp. ARC (Eddie *et al*., 2008) and *C.* sp. ICE-L, ICE-W (Liu *et al*., 2006), ICE-MDV (ref) and KNF0022 (Jung *et al*., 2016). These patterns raise the possibility of a ‘Polar clade’ within the Chlamydomonadales, enriched in psychrophilic species.

These close phylogenetic relationships despite geographical distance raise several possibilities that could explain the ancestry of psychrophilic green algae. For example, the Arctic-Antarctic split within this group could result from ancient climate history rather than modern geographical connectivity. Snow- and ice-associated *Chlamydomonas* are often capable of forming life stages resilient to desiccation, freezing, and high UV exposure (Hoham & Remias, 2020) that could facilitate long-distance atmospheric or oceanic dispersal, particularly during colder geological periods (e.g., the Pleistocene glaciation, (Clark *et al*., 2009). The extreme conditions and prolonged isolation between the Arctic and Antarctic could favor a narrow set of pre-adapted lineages and result in a closely related but geographically isolated species that occupy comparable polar niches (Dorrell *et al*., 2023). Similar theories have been suggested for the dispersal of cold-adapted Trebouxiophyceae, diatoms and foraminifers (Darling *et al*., 2000; Vyverman *et al*., 2007; Hodač *et al*., 2016; Segawa *et al*., 2018, 2023) and such events could have shaped modern polar *Chlamydomonas* diversity as well. The genus *Chlamydomonas* could be especially predisposed to bipolar evolutionary patterns: species within this group occupy a wide range of aquatic habitats, including snow, glacier ice, fresh waters, brackish and marine environments, and have repeatedly developed traits for cold tolerance, osmotic flexibility, and stress resilience (Leliaert *et al*., 2012; Hoham & Remias, 2020). Molecular clock analyses are currently limited by sparse taxon sampling and high likelihood of unsampled diversity (Rappaport & Oliverio, 2023), and broader genomic representation will enable divergence-time estimates across this group in the future.

### Less extreme conditions promote algal growth and phenotypic change

Light and salinity are major environmental drivers for microbial growth in polar environments (Li *et al*., 2019; Duncan *et al*., 2024a,b; Ahmed *et al*., 2025). All three psychrophiles examined here exhibited the slowest growth rates when acclimated to shading (Fig. 2a), conditions most often associated with ice- or snow-covered environments (Stibal *et al*., 2007; Smoot & Hopcroft, 2017; Patriarche *et al*., 2021). Conversely, their population increased the fastest when acclimated to less extreme conditions. Light was a major driver of high growth rates, and particularly when combined with lower salinity (Fig. 2a). These results are consistent with published lab-based studies on axenic algal cultures (Takizawa *et al*., 2009; Pocock *et al*., 2011; Lee *et al*., 2018; Suzuki *et al*., 2019; Morales-Sánchez *et al*., 2020a; Stahl-Rommel *et al*., 2021), but it is also reminiscent of recent ecological observations in polar environments. The loss of the ice- and snow-cover and freshening of marine habitats due to warming, and the consequent increase in greening and primary productivity, have been well documented in both Arctic (Arrigo *et al*., 2012; Myers-Smith *et al*., 2020b; Castellani *et al*., 2022; Manizza *et al*., 2023; Wolf *et al*., 2024) and Antarctic habitats (van Leeuwe *et al*., 2018; Del Castillo *et al*., 2019; Gray *et al*., 2020; Roland *et al*., 2024).

In addition to high growth rates, we also observed higher palmelloid content in all three psychrophiles acclimated to higher light, regardless of the salinity (Fig. 2c). Palmelloid formation may be a general stress response in green algae and has been observed upon expose to pollutants, salinity, nutrient deficiency, and grazers (Lurling & Beekman, 2006; Ratcliff *et al*., 2013; Khona *et al*., 2016; Cheloni & Slaveykova, 2021; Pascual *et al*., 2025) but there is evidence that this morphological structure may be photoprotective. The mesophilic *C. reinhardtii* utilized the formation of palmelloids and aggregates to shield internal cells from excessive light and promote increases in antioxidant content (Suwannachuen *et al*., 2023). The Antarctic *C. priscui* grown at suboptimal temperatures (16°C) induced the formation of palmelloids with modified organization of the photosynthetic apparatus and enhanced capacity for quenching of excess absorbed light (Szyszka-Mroz *et al*., 2022; Hüner *et al*., 2023). Similar morphology has been reported alpine algal biofilms, suggesting a protective role against excessive light and UV exposure in snow communities (Karsten & Holzinger, 2014). We observed a light-driven increase in palmelloid content in all three species, regardless of their native habitat (Fig. 2c), suggesting that a change in phenotype from a motile single cell to a non-motile colony may be a general photoprotective response. However, it must be noted that the light intensities used in our work are not likely to induce severe photodamage. For instance, *C. priscui* exhibited robust growth and enhanced phototolerance at 250 μmol m^-2^ s^-1^, light intensities equivalent to those measured in shallow open moats in Antarctic lakes (Popson *et al*., 2024), and algae growing at the snow-air interface routinely experience very high irradiances (1,000-2,000 μmol m^-2^ s^-1^) (Hoham & Remias, 2020). Thus, a comprehensive assessment of palmelloid formation in psychrophilic algae requires testing at light intensities near the upper permissible range, where the photoprotective role of this phenotype is most likely to become apparent.

### Resilience to temperature stress is dependent on acclimation to light and salinity

A highlight of this study is the observation that all three psychrophiles acclimated to less extreme conditions (higher light intensity, lower salinity) exhibit rapid growth but also the highest sensitivity to temperature stress. All three species acclimated to these “relaxed” conditions lost viability and chlorophyll within 3-4 days when challenged with non-permissive temperatures (Fig. 3). High growth rates also correlate to a significant and rapid deterioration of F_V_/F_M_ (Fig. 4a-c), electron transport rates (Fig. d-f), and photochemical yields (Fig. 5a-c) within 24-48 hrs of temperature stress. In contrast, slow-growing algal cultures acclimated to high salinity and low light intensity were much more robust. Complete loss of viability and chlorophyll required ∼15 days of temperature stress (Fig. 3), and these cultures maintained the efficiency of their photosynthetic apparatus (Fig. 4, Fig. 5) over 48 hrs at increased temperatures. *C. priscui* is the only psychrophilic alga that has been examined in the context of temperature stress, but all previous studies were done on cultures acclimated to low salinity and higher light. This alga constitutively accumulates stress-related metabolites and proteins (e.g., heat shock proteins, antioxidants) during low temperature growth, like as an adaptation to the extreme conditions in Lake Bonney, but fails to further increase their levels during short-term acute temperature stress (Cvetkovska *et al*., 2022b). Longer exposure to heat leads to rapid cell death, loss of Y(II) and consequent increases in Y(NO) and Y(NPQ) (Possmayer *et al*., 2011). These studies suggested that adaptations that may confer a benefit for life at the extreme, may also limit the ability of psychrophilic algae to respond to environmental disturbances (Cvetkovska *et al*., 2022a), but here we demonstrate that these responses are highly dependent on algal acclimation to different light and salinity levels. Our work paves the way towards detailed examination of the molecular basis that underlies the sensitivity of polar algae to environmental change and the roles of salinity, light, and temperature in stress resilience.

While these general trends hold true in all three psychrophiles examined here, we also show that the interaction between light, salinity, and temperature is complex. For instance, our data suggests that salinity is the main driver of temperature sensitivity in *C. priscui*, while light intensity more strongly affects stress resilience in *C. klinobasis*. The temperature sensitivity in *C. malina* appears to be governed by both light and salinity (Fig. 4, Table S3). These patterns could reflect the species native habitats. *C. priscui* is endemic to the perennially ice-covered deep waters of Lake Bonney. This is an extremely light-limited environment with dramatic increases in conductivity within a sharp chemocline at ∼17-20m, which coincides with the depth at which *C. priscui* was isolated (Neale & Priscu, 1995; Patriarche *et al*., 2021). The freshwater alga *C. klinobasis* was isolated from ephemeral snowfields on Svalbard Island (Hulatt *et al*., 2017), a dynamic oligotrophic environment that experiences significant light variability: extremely high irradiation on the surface with rapid light attenuation within the deeper snow layers (Stibal *et al*., 2007). Finally, the Arctic marine alga *C. malina* is native to the Beaufort Sea (Balzano *et al*., 2012), a highly dynamic habitat characterized by a thick ice cover in the winter, and thinner ice packs and open waters during the summer. In addition, this environment receives significant freshwater and nutrient inputs from the surrounding land masses (Tremblay *et al*., 2008; Rheinlænder *et al*., 2024). Thus, the responses of these psychrophiles to temperature stress are likely shaped by life at cold, but otherwise very different, native habitats.

Cross-tolerance between salinity and heat has been reported in mesophiles, including a similar response in *Dunaliella* sp., which was progressively more resilient to heat stress when cultured at increasing salinities (Henley *et al*., 2002). Similarly, exposure to moderate salinities lead to improved resilience in plants (Rivero *et al*., 2014; Pushpavalli *et al*., 2020) and coral endosymbionts (Gegner *et al*., 2017), suggesting a conserved response. The mechanisms behind the protective role of salinity is not fully understood, but moderate salinity stress enhances the accumulation of protective metabolites (e.g., proline, glycine, sucrose), antioxidants (e.g., ascorbate) and cytosolic Ca^2+^-related signalling cascades (Rivero *et al*., 2014; Shaar-Moshe *et al*., 2017; Ji *et al*., 2018; Guo *et al*., 2022) likely priming for broader stress resilience. Indeed, high salinity-acclimated *C. priscui* cultures accumulate high levels of antioxidants, proline, and exhibit enhanced CEF around PSI (Kalra *et al*., 2020, 2023; Stahl-Rommel *et al*., 2021). Similar mechanisms likely confer a broad-spectrum resilience to temperature stress in all psychrophiles examined here (Fig. 4, Fig. 5).

The interaction between light and elevated temperature is considered antagonistic in photosynthetic organisms, as higher irradiances can exacerbate heat stress by increasing PSII excitation pressure, accelerate photoinhibition, and increase chloroplast-derived ROS accumulation (Allakhverdiev *et al*., 2008; Gollan & Aro, 2020; Zahra *et al*., 2023). Possmayer et al. (2011) demonstrated that a shift to a non-permissive temperature under growth irradiance (100 μmol m^-2^ s^-1^) led to significantly increased kinetics of cell death in *C. priscui*, compared to equivalent temperature stress applied to cultures in the dark. While this alga is capable of long-term acclimation to higher irradiance (100-250 μmol m^-2^ s^-1^) due to its robust antioxidant system, high NPQ capacity, and the protective role of CEF (Szyszka *et al*., 2007; Popson *et al*., 2024; Poirier *et al*., 2025), these mechanisms are likely insufficient to counteract the multifactorial effect of combined higher irradiance and temperature stress (Fig. 4a,d, Fig. 5a). While similar processes are likely to operate in *C. malina* (Fig. 4b, e, Fig. 5b), the effects of light may differ in *C. klinobasis* (Fig. 4c, f, Fig. 5c). Among the psychrophiles examined in this work, the snow alga *C. klinobasis* displayed the highest temperature stress resilience at the level of photosynthesis and maintained photosynthetic efficiency over 48 hrs in all conditions (Fig. 5). Snow algae have well characterized protection against extreme irradiation due to a robust capacity for the synthesis of astaxanthin and other carotenoids. These pigments have photoprotective and antioxidant function, act as metabolic sinks, and may physically shade organelles from light damage (Hoham & Remias, 2020). Such mechanisms may contribute to the stress resilience of *C. klinobasis* observed here.

### Implications for polar photosynthetic life during climate change

Our work is the first to directly correlate the acclimation to different light and salinity levels with temperature stress resilience in psychrophiles (Fig. 6). The effects of temperature, light, and salinity of the physiology of psychrophilic algae have been examined in a handful of studies (Morales-Sánchez *et al*., 2020a; Young & Schmidt, 2020; Morales-Sánchez *et al*., 2020b; Cvetkovska *et al*., 2022a; Hüner *et al*., 2023; Morgan-Kiss *et al*., 2024), but previous work has largely centered on a single environmental variable. Light and salinity are the main drivers of algal diversity and productivity in polar environments, and the ones that rapidly change with the loss of ice cover in the warming polar regions (Castendyk *et al*., 2016; Lehnherr *et al*., 2018; Li *et al*., 2019; Obryk *et al*., 2019; Meiners *et al*., 2025). Cold-water organisms will be further challenged with short-term but severe episodes of anomalous weather patterns, including more frequent extreme heat waves (Huang *et al*., 2021) with recorded temperatures rising ∼40°C above the yearly average (Constable, 2022; Blanchard-Wrigglesworth *et al*., 2023).

**Figure 6:**
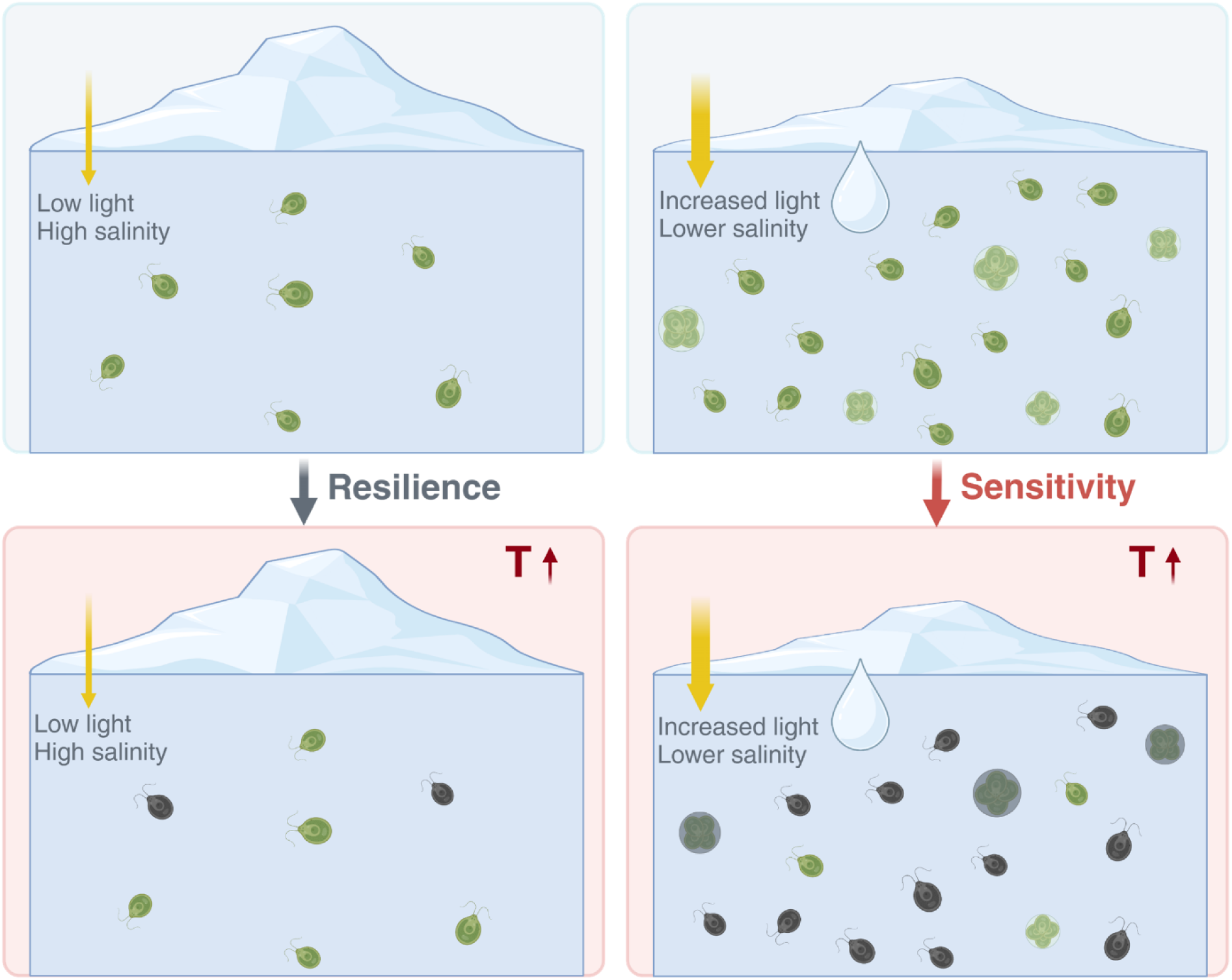
A model representing the responses of psychrophilic *Chlamydomonas* to a changing environment. Green algae acclimated to higher salinities and lower light intensities exhibit slow growth (top left) but maintain high resilience to subsequent lethal heat stress (bottom left). Conversely, green algae acclimated to increased light intensities and decreased salinities, conditions that may result from ice and snow melt, grow rapidly (top right) but are sensitive to lethal heat stress and display rapid cellular death (bottom right)

Our results suggest that fast-growing psychrophilic algae acclimated to lower salinities and increasing light intensities are at particular risk (Fig. 6). This is likely to have major implications in polar habitats, and particularly in ice-covered Antarctic lakes and snow fields dominated by psychrophilic *Chlamydomonas* (Li *et al*., 2019; Hoham & Remias, 2020; Morgan-Kiss *et al*., 2024). Indeed, the once highly abundant *C. priscui* (Neale & Priscu, 1995) is only rarely detected in the Antarctic Lake Bonney in recent surveys (Li *et al*., 2019; Morgan-Kiss *et al*., 2024) and may be one of the casualties of climate change. Our work was done on axenic cultures but natural microbial communities are complex and many species are psychrotolerant rather than psychrophilic (Hüner *et al*., 2022). Rising temperatures may give a competitive advantage to psychrotolerant algae that survive in the cold but have growth maxima at temperatures ≥20°C; thus, fine-grained analyses on the physiology of psychrophilic algae faced with changing conditions must be complemented with long-term observations of algal population dynamics in the polar regions.

Natural habitats are characterized by co-occurring or sequential stressors, and a conceptual framework known as the multifactorial stress combination (MFSC) principle has been proposed to address the effect of multiple conditions on an organism (Zandalinas & Mittler, 2022). MFSC responses have been explored in plants (Zandalinas *et al*., 2021) and soil microbial communities (Rillig *et al*., 2019), but how multifactorial and dynamic conditions affect psychrophilic algae is still elusive. Our work provides the first advancement in mapping the responses of polar algae to a changing environment and opens to doors to future studies on the acclimation and stress resilience in these important primary producers.

## Acknowledgements

This project was supported by Natural Sciences and Engineering Research Council of Canada Discovery Grants (NSERC DG) awarded to M.C. The authors are grateful for the support from the Canada Foundation for Innovation (CFI) and University of Ottawa start-up funding. P.O. was supported by Ontario Graduate Scholarship (OGS) and NSERC Graduate Scholarship. The authors wish to thank Rhys Thushingham for their help in algal culturing, and Marc Possmayer for helpful discussion on algal phylogenetic analysis.

## Autor Contributions

PO and MC designed the study and interpreted the data. PO performed all physiology experiments and analyzed the data. SC and NB assisted with algal culturing and preformed the initial growth characterization of *C. malina* and *C. klinobasis*. AS and LK contributed to sequence data collection and phylogenetic analyses. PO and MC wrote the original draft. All authors edited and approved the manuscript.

## Data availability

All data is available from the corresponding author on request. Sequences for RbcL from *C. malina* (PZ227531) and *C. klinobasis* (PZ227532) are deposited in GenBank.

## Supporting information

**Figure S1:** Phylogenetic tree of 63 *RbcL* nucleotide sequences inferred from using maximum likelihood analysis is shown to scale (substitutions per site), with branch values representing bootstrap support. GenBank accession numbers are provided in brackets beside the organism name. The species considered in this study are highlighted in blue (*Chlamydomonas priscui*), green (*Chlamydomonas malina*) and mauve (*Chlamydomonas klinobasis*).

**Figure S2:** Phylogenetic tree of 111 18S *rRNA* nucleotide sequences inferred from using maximum likelihood analysis is shown to scale (substitutions per site), with branch values representing bootstrap support. GenBank accession numbers are provided in brackets beside the organism name. The species considered in this study are highlighted in blue (*Chlamydomonas priscui*), green (*Chlamydomonas malina*) and mauve (*Chlamydomonas klinobasis*).

**Figure S3:** Optimizing the growth and stress conditions in *C. klinobasis* and *C. malina*. (**a**) Optical density measured at a range of salinities (0.43-70 mM NaCl) in *C. klinobasis* cultures. (**b**) Optical densities measured at a range of salinities (10-600 mM) in *C. malina* cultures. In both (a) and (b), algal cultures cultured at 4°C. (c) Kinetics of cell death in *C. klinobasis* exposed to non-permissive temperatures (21°C, 22°C, 23°C). (d) Kinetics of cell death in *C. malina* exposed to non-permissive temperatures (22°C, 23°C, 24°C).

**Table S1:** A comparison of the growth rates in *C. priscui* (P), *C. malina* (M), and *C. klinobasis* (K) cultured at 4°C and acclimated to high salinity (HS), low salinity (LS), high light (HL) or low light (LL). Data are means ± SD of at least three biological replicates. The lack of statistical difference (p > 0.05) between the growth rates of the three species was confirmed with a one-way ANOVA with a Tukey’s post-hoc test preformed within each treatment.

**Table S2:** A statistical summary of two-way ANOVA analyses for the interaction effects between the experimental variables (salinity, light intensity) and the measured morphological and physiological parameters in *C. priscui*, *C. malina* and *C. klinobasis.* The numbers represent p-values, and significant interactions (*p* < 0.05) are bolded.

**Table S3:** A statistical summary of three-way ANOVA analyses for the interaction effects between the experimental variables (salinity, light intensity, time at non-permissive temperature) and the measured viability and physiological parameters in *C. priscui*, *C. malina* and *C. klinobasis.* The numbers represent p-values, and significant interactions (*p* < 0.05) are bolded.

**Table S4:** A statistical summary of three-way ANOVA analyses followed by Dunnett’s post hoc test for the energy partitioning parameters measured in *C. prisui*, *C. malina* and *C. klinobasis*. The total absorbed light is distributed between photochemical yield of PSII Y(II), non-photochemical quenching Y(NPQ), and non-regulated dissipation Y(NO). In all cases, algae were cultured at 4°C and acclimated to high salinity (HS), low salinity (LS), high light (HL) or low light (LL). Temperature stress was initiated at the mid-exponential growth phase (OD750 = 0.3-0.4) and corresponds to t = 0 hours. Data are means ± SD of at least three biological replicates. Statistical significance is indicated by an asterisk and bolded (* = < 0.05, ** = < 0.01, *** = < 0.001, **** = < 0.0001).

## References

Ahmed F, Leu E, Juhl AR, Campbell K, Dilliplaine KB, Assmy P, Niemi A, Gradinger R, Alou-Font E, Torres-Valdés S, et al. 2025. A pan-Arctic perspective on the influence of ice algae on sea-ice nutrient concentrations. Elementa: Science of the Anthropocene 13: 00059.

Allakhverdiev SI, Kreslavski VD, Klimov VV, Los DA, Carpentier R, Mohanty P. 2008. Heat stress: an overview of molecular responses in photosynthesis. Photosynthesis Research 98: 541.

Anderson LG, and Macdonald RW. 2015. Observing the Arctic Ocean carbon cycle in a changing environment. Polar Research 34: 26891.

Arrigo KR, Perovich DK, Pickart RS, Brown ZW, van Dijken GL, Lowry KE, Mills MM, Palmer MA, Balch WM, Bahr F, et al. 2012. Massive Phytoplankton Blooms Under Arctic Sea Ice. Science 336: 1408–1408.

Balzano S, Gourvil P, Siano R, Chanoine M, Marie D, Lessard S, Sarno D, Vaulot D. 2012. Diversity of cultured photosynthetic flagellates in the northeast Pacific and Arctic Oceans in summer. Biogeosciences 9: 4553–4571.

Bax N, Sands CJ, Gogarty B, Downey RV, Moreau CVE, Moreno B, Held C, Paulsen ML, McGee J, Haward M, et al. 2021. Perspective: Increasing blue carbon around Antarctica is an ecosystem service of considerable societal and economic value worth protecting. Global Change Biology 27: 5–12.

Bers AV, Momo F, Schloss IR, Abele D. 2013. Analysis of trends and sudden changes in long-term environmental data from King George Island (Antarctica): relationships between global climatic oscillations and local system response. Climatic Change 116: 789–803.

Blanchard-Wrigglesworth E, Cox T, Espinosa ZI, Donohoe A. 2023. The Largest Ever Recorded Heatwave—Characteristics and Attribution of the Antarctic Heatwave of March 2022. Geophysical Research Letters 50: e2023GL104910.

Bronselaer B, Winton M, Griffies SM, Hurlin WJ, Rodgers KB, Sergienko OV, Stouffer RJ, Russell JL. 2018. Change in future climate due to Antarctic meltwater. Nature 564: 53–58.

Buchheim MA, Michalopulos EA, Buchheim JA. 2001. PHYLOGENY OF THE CHLOROPHYCEAE WITH SPECIAL REFERENCE TO THE SPHAEROPLEALES: A STUDY OF 18S AND 26S rDNA DATA. Journal of Phycology 37: 819–835.

Cárdenas CA, González-Aravena M, Santibañez PA. 2018. The importance of local settings: within-year variability in seawater temperature at South Bay, Western Antarctic Peninsula. PeerJ 6: e4289.

Castellani G, Veyssière G, Karcher M, Stroeve J, Banas SN, Bouman AH, Brierley SA, Connan S, Cottier F, Große F, et al. 2022. Shine a light: Under-ice light and its ecological implications in a changing Arctic Ocean. Ambio 51: 307–317.

Castendyk DN, Obryk MK, Leidman SZ, Gooseff M, Hawes I. 2016. Lake Vanda: A sentinel for climate change in the McMurdo Sound Region of Antarctica. Global and Planetary Change 144: 213–227.

Cheloni G, Slaveykova VI. 2021. Morphological plasticity in *Chlamydomonas reinhardtii* and acclimation to micropollutant stress. Aquatic Toxicology 231: 105711.

Clark PU, Dyke AS, Shakun JD, Carlson AE, Clark J, Wohlfarth B, Mitrovica JX, Hostetler SW, McCabe AM. 2009. The Last Glacial Maximum. Science 325: 710–714.

Constable AJ. 2022. Imperatives for integrated science and policy in managing greenhouse gas risks to the Southern Polar Region. Global Change Biology 28: 4489–4492.

Cook G, Teufel A, Kalra I, Li W, Wang X, Priscu J, Morgan-Kiss R. 2019. The Antarctic psychrophiles Chlamydomonas spp. UWO241 and ICE-MDV exhibit differential restructuring of photosystem I in response to iron. Photosynthesis Research 141: 209–228.

Criscuolo A, Gribaldo S. 2010. BMGE (Block Mapping and Gathering with Entropy): a new software for selection of phylogenetic informative regions from multiple sequence alignments. BMC Evolutionary Biology 10: 210.

Cvetkovska M, Hüner NPA, Smith DR. 2017. Chilling out: the evolution and diversification of psychrophilic algae with a focus on Chlamydomonadales. Polar Biology 40: 1169–1184.

Cvetkovska M, Orgnero S, Hüner NPA, Smith DR. 2019. The enigmatic loss of light-independent chlorophyll biosynthesis from an Antarctic green alga in a light-limited environment. New Phytologist 222: 651–656.

Cvetkovska M, Vakulenko G, Smith DR, Zhang X, Hüner NPA. 2022a. Temperature stress in psychrophilic green microalgae: Minireview. Physiologia Plantarum 174: e13811.

Cvetkovska M, Zhang X, Vakulenko G, Benzaquen S, Szyszka-Mroz B, Malczewski N, Smith DR, Hüner NPA. 2022b. A constitutive stress response is a result of low temperature growth in the Antarctic green alga *Chlamydomonas* sp. UWO241. Plant, Cell & Environment 45: 156–177.

Dalpadado P, Arrigo KR, van Dijken GL, Skjoldal HR, Bagøien E, Dolgov AV, Prokopchuk IP, Sperfeld E. 2020. Climate effects on temporal and spatial dynamics of phytoplankton and zooplankton in the Barents Sea. Progress in Oceanography 185: 102320.

Darling KF, Wade CM, Stewart IA, Kroon D, Dingle R, Brown AJL. 2000. Molecular evidence for genetic mixing of Arctic and Antarctic subpolar populations of planktonic foraminifers. Nature 405: 43–47.

Davey MP, Norman L, Sterk P, Huete-Ortega M, Bunbury F, Loh BKW, Stockton S, Peck LS, Convey P, Newsham KK, et al. 2019. Snow algae communities in Antarctica: metabolic and taxonomic composition. New Phytologist 222: 1242–1255.

Del Castillo CE, Signorini S, Karaköylü EM, Rivero-Calle S. 2019. Is the Southern Ocean getting greener? Geophysical research letters 46: 6034–6040.

Demchenko E, Mikhailyuk T, Coleman AW, Pröschold T. 2012. Generic and species concepts in Microglena (previously the Chlamydomonas monadina group) revised using an integrative approach. European Journal of Phycology 47: 264–290.

Dorrell RG, Kuo A, Füssy Z, Richardson EH, Salamov A, Zarevski N, Freyria NJ, Ibarbalz FM, Jenkins J, Pierella Karlusich JJ, et al. 2023. Convergent evolution and horizontal gene transfer in Arctic Ocean microalgae. Life Science Alliance 6: e202201833.

Duncan RJ, Nielsen D, Søreide JE, Varpe Ø, Tobin MJ, Pitusi V, Heraud P, Petrou K. 2024a. Biomolecular profiles of Arctic sea-ice diatoms highlight the role of under-ice light in cellular energy allocation. ISME Communications 4: ycad010.

Duncan RJ, Søreide JE, Varpe Ø, Wiktor J, Pitusi V, Runge E, Petrou K. 2024b. Spatio-temporal dynamics in microalgal communities in Arctic land-fast sea ice. Progress in Oceanography 224: 103248.

Durack PJ, Wijffels SE. 2010. Fifty-Year Trends in Global Ocean Salinities and Their Relationship to Broad-Scale Warming.

Eddie B, Krembs C, Neuer S. 2008. Characterization and growth response to temperature and salinity of psychrophilic, halotolerant Chlamydomonas sp. ARC isolated from Chukchi Sea ice. Marine Ecology Progress Series 354: 107–117.

Feron S, Cordero RR, Damiani A, Malhotra A, Seckmeyer G, Llanillo P. 2021. Warming events projected to become more frequent and last longer across Antarctica. Scientific Reports 11: 19564.

Gegner HM, Ziegler M, Rädecker N, Buitrago-López C, Aranda M, Voolstra CR. 2017. High salinity conveys thermotolerance in the coral model Aiptasia. Biology Open 6: 1943–1948.

Gérikas Ribeiro C, dos Santos AL, Gourvil P, Le Gall F, Marie D, Tragin M, Probert I, Vaulot D. 2020. Culturable diversity of Arctic phytoplankton during pack ice melting (JW Deming and C Michel, Eds). Elementa: Science of the Anthropocene 8: 6.

Gollan PJ, Aro E-M. 2020. Photosynthetic signalling during high light stress and recovery: targets and dynamics. Philosophical Transactions of the Royal Society B: Biological Sciences 375: 20190406.

Gray A, Krolikowski M, Fretwell P, Convey P, Peck LS, Mendelova M, Smith AG, Davey MP. 2020. Remote sensing reveals Antarctic green snow algae as important terrestrial carbon sink. Nature Communications 11: 2527.

Guindon S, Dufayard J-F, Lefort V, Anisimova M, Hordijk W, Gascuel O. 2010. New Algorithms and Methods to Estimate Maximum-Likelihood Phylogenies: Assessing the Performance of PhyML 3.0. Systematic Biology 59: 307–321.

Guo M, Wang X-S, Guo H-D, Bai S-Y, Khan A, Wang X-M, Gao Y-M, Li J-S. 2022. Tomato salt tolerance mechanisms and their potential applications for fighting salinity: A review. Frontiers in Plant Science 13.

Hadi SIIA, Santana H, Brunale PPM, Gomes TG, Oliveira MD, Matthiensen A, Oliveira MEC, Silva FCP, Brasil BSAF. 2016. DNA Barcoding Green Microalgae Isolated from Neotropical Inland Waters. PLOS ONE 11: e0149284.

Henley WJ, Major KM, Hironaka JL. 2002. Response to Salinity and Heat Stress in Two Halotolerant Chlorophyte Algae1. Journal of Phycology 38: 757–766.

Hill VJ, Light B, Steele M, Zimmerman RC. 2018. Light Availability and Phytoplankton Growth Beneath Arctic Sea Ice: Integrating Observations and Modeling. Journal of Geophysical Research: Oceans 123: 3651–3667.

Hodač L, Hallmann C, Spitzer K, Elster J, Faßhauer F, Brinkmann N, Lepka D, Diwan V, Friedl T. 2016. Widespread green algae Chlorella and Stichococcus exhibit polar-temperate and tropical-temperate biogeography. FEMS microbiology ecology 92: fiw122.

Hoham RW, Remias D. 2020. Snow and Glacial Algae: A Review1. Journal of Phycology 56: 264–282.

Huang B, Wang Z, Yin X, Arguez A, Graham G, Liu C, Smith T, Zhang H-M. 2021. Prolonged Marine Heatwaves in the Arctic: 1982−2020. Geophysical Research Letters 48: e2021GL095590.

Hui C, Schmollinger S, Glaesener AG. 2023. Chapter 11 - Growth techniques. In: Goodenough U, ed. The Chlamydomonas Sourcebook (Third Edition). Academic Press, 287–314.

Hulatt CJ, Berecz O, Egeland ES, Wijffels RH, Kiron V. 2017. Polar snow algae as a valuable source of lipids? Bioresource Technology 235: 338–347.

Hüner NPA, Ivanov AG, Szyszka-Mroz B, Savitch LV, Smith DR, Kata V. 2024. Photostasis and photosynthetic adaptation to polar life. Photosynthesis Research 161: 51–64.

Hüner NPA, Smith DR, Cvetkovska M, Zhang X, Ivanov AG, Szyszka-Mroz B, Kalra I, Morgan-Kiss R. 2022. Photosynthetic adaptation to polar life: Energy balance, photoprotection and genetic redundancy. Journal of Plant Physiology 268: 153557.

Hüner NPA, Szyszka-Mroz B, Ivanov AG, Kata V, Lye H, Smith DR. 2023. Photosynthetic adaptation and multicellularity in the Antarctic psychrophile, *Chlamydomonas priscuii*. Algal Research 74: 103220.

Jeffrey SW, Humphrey GF. 1975. New spectrophotometric equations for determining chlorophylls *a*, *b*, *c*1 and *c*2 in higher plants, algae and natural phytoplankton. Biochemie und Physiologie der Pflanzen 167: 191–194.

Ji X, Cheng J, Gong D, Zhao X, Qi Y, Su Y, Ma W. 2018. The effect of NaCl stress on photosynthetic efficiency and lipid production in freshwater microalga—*Scenedesmus obliquus* XJ002. Science of The Total Environment 633: 593–599.

Jung W, Kim EJ, Lim S, Sim H, Han SJ, Kim S, Kang S-H, Choi H-G, Jung W, Kim EJ, et al. 2016. Cellular growth and fatty acid content of Arctic chlamydomonadalean. Algae 31: 61–71.

Kalra I, Wang X, Cvetkovska M, Jeong J, McHargue W, Zhang R, Hüner NPA, Yuan JS, Morgan-Kiss RM. 2020. Chlamydomonas sp. UWO 241 exhibits high cyclic electron flow and rewired metabolism under high salinity. Plant Physiology: pp.01280.2019.

Kalra I, Wang X, Zhang R, Morgan-Kiss R. 2023. High salt-induced PSI-supercomplex is associated with high CEF and attenuation of state transitions. Photosynthesis Research 157: 65–84.

Karsten U, Holzinger A. 2014. Green algae in alpine biological soil crust communities: acclimation strategies against ultraviolet radiation and dehydration. Biodiversity and Conservation 23: 1845–1858.

Katoh K, Standley DM. 2013. MAFFT Multiple Sequence Alignment Software Version 7: Improvements in Performance and Usability. Molecular Biology and Evolution 30: 772–780.

Khona DK, Shirolikar SM, Gawde KK, Hom E, Deodhar MA, D’Souza JS. 2016. Characterization of salt stress-induced palmelloids in the green alga, *Chlamydomonas reinhardtii*. Algal Research 16: 434–448.

Kramer DM, Johnson G, Kiirats O, Edwards GE. 2004. New Fluorescence Parameters for the Determination of QA Redox State and Excitation Energy Fluxes. Photosynthesis Research 79: 209–218.

Lee K-K, Lim P-E, Poong S-W, Wong C-Y, Phang S-M, Beardall J. 2018. Growth and photosynthesis of Chlorella strains from polar, temperate and tropical freshwater environments under temperature stress. Journal of Oceanology and Limnology 36: 1266–1279.

van Leeuwe MA, Tedesco L, Arrigo KR, Assmy P, Campbell K, Meiners KM, Rintala J-M, Selz V, Thomas DN, Stefels J. 2018. Microalgal community structure and primary production in Arctic and Antarctic sea ice: A synthesis (JW Deming, Ed.). Elementa: Science of the Anthropocene 6: 4.

Lefort V, Longueville J-E, Gascuel O. 2017. SMS: Smart Model Selection in PhyML. Molecular Biology and Evolution 34: 2422–2424.

Lehnherr I, St. Louis VL, Sharp M, Gardner AS, Smol JP, Schiff SL, Muir DCG, Mortimer CA, Michelutti N, Tarnocai C, et al. 2018. The world’s largest High Arctic lake responds rapidly to climate warming. Nature Communications 9: 1290.

Leliaert F, Smith DR, Moreau H, Herron MD, Verbruggen H, Delwiche CF, De Clerck O. 2012. Phylogeny and Molecular Evolution of the Green Algae. Critical Reviews in Plant Sciences 31: 1–46.

Lemieux C, Vincent AT, Labarre A, Otis C, Turmel M. 2015. Chloroplast phylogenomic analysis of chlorophyte green algae identifies a novel lineage sister to the Sphaeropleales (Chlorophyceae). BMC Evolutionary Biology 15: 264.

Lemoine F, Correia D, Lefort V, Doppelt-Azeroual O, Mareuil F, Cohen-Boulakia S, Gascuel O. 2019. NGPhylogeny.fr: new generation phylogenetic services for non-specialists. Nucleic Acids Research 47: W260–W265.

Letunic I, Bork P. 2021. Interactive Tree Of Life (iTOL) v5: an online tool for phylogenetic tree display and annotation. Nucleic Acids Research 49: W293–W296.

Li W, Dolhi-Binder J, Cariani Z, Morgan-Kiss R. 2019. Drivers of protistan community autotrophy and heterotrophy in chemically stratified Antarctic lakes. Aquatic Microbial Ecology 82: 225–239.

Liu C, Huang X, Wang X, Zhang X, Li G. 2006. Phylogenetic studies on two strains of Antarctic ice algae based on morphological and molecular characteristics. Phycologia 45: 190–198.

Lodeyro AF, Krapp AR, Carrillo N. 2021. Photosynthesis and chloroplast redox signaling in the age of global warming: stress tolerance, acclimation, and developmental plasticity. Journal of Experimental Botany 72: 5919–5937.

Lurling M, Beekman W. 2006. Palmelloids formation in Chlamydomonas reinhardtii : defence against rotifer predators? Annales de Limnologie - International Journal of Limnology 42: 65–72.

Malapascua J, Jerez C, Sergejevová M, Figueroa F, Masojídek J. 2014. Photosynthesis monitoring to optimize growth of microalgal mass cultures: application of chlorophyll fluorescence techniques. Aquatic Biology 22: 123–140.

Manizza M, Carroll D, Menemenlis D, Zhang H, Miller CE. 2023. Modeling the Recent Changes of Phytoplankton Blooms Dynamics in the Arctic Ocean. Journal of Geophysical Research: Oceans 128: e2022JC019152.

Meiners KM, Lannuzel D, Leu E, Brown KA. 2025. Nutrients and organic matter dynamics in sea ice. In: Sea Ice. John Wiley & Sons, Ltd, 575–622.

Morales-Sánchez D, Schulze PSC, Kiron V, Wijffels RH. 2020a. Temperature-Dependent Lipid Accumulation in the Polar Marine Microalga Chlamydomonas malina RCC2488. Frontiers in Plant Science 11.

Morales-Sánchez D, Schulze PSC, Kiron V, Wijffels RH. 2020b. Production of carbohydrates, lipids and polyunsaturated fatty acids (PUFA) by the polar marine microalga *Chlamydomonas malina* RCC2488. Algal Research 50: 102016.

Morgan-Kiss, R.M., Guiry, M.D. 2024. Validation of “*Chlamydomonas priscui*” (“*Chlamydomonas* sp. UWO241”; Chlamydomonadaceae, Chlorophyceae), a psychrophilic organism from the Antarctic widely used in research on cold adaptation of photosynthesis. Notulae Algarum 346: 1–2.

Morgan-Kiss RM, Popson D, Pereira R, Dolhi-Binder J, Teufel A, Li W, Kalra I, Sherwell S, Reynebeau E, Takacs-Vesbach C. 2024. Sentinel protist taxa of the McMurdo Dry Valley lakes, Antarctica: a review. Frontiers in Ecology and Evolution 12: 1323472.

Morita RY. 1975. Psychrophilic bacteria. Bacteriological Reviews 39: 144–167.

Myers-Smith IH, Kerby JT, Phoenix GK, Bjerke JW, Epstein HE, Assmann JJ, John C, Andreu-Hayles L, Angers-Blondin S, Beck P, et al. 2020a. Complexity revealed in the greening of the Arctic. NATURE CLIMATE CHANGE.

Myers-Smith IH, Kerby JT, Phoenix GK, Bjerke JW, Epstein HE, Assmann JJ, John C, Andreu-Hayles L, Angers-Blondin S, Beck PSA, et al. 2020b. Complexity revealed in the greening of the Arctic. Nature Climate Change 10: 106–117.

Nakada T, Misawa K, Nozaki H. 2008. Molecular systematics of Volvocales (Chlorophyceae, Chlorophyta) based on exhaustive 18S *r*RNA phylogenetic analyses. Molecular Phylogenetics and Evolution 48: 281–291.

Nakada T, Tomita M, Wu J-T, Nozaki H. 2016. Taxonomic revision of Chlamydomonas subg. Amphichloris (Volvocales, Chlorophyceae), with resurrection of the genus Dangeardinia and descriptions of Ixipapillifera gen. nov. and Rhysamphichloris gen. nov. Journal of Phycology 52: 283–304.

Neale PJ, Priscu JC. 1995. The Photosynthetic Apparatus of Phytoplankton from a Perennially Ice-Covered Antarctic Lake: Acclimation to an Extreme Shade Environment. Plant and Cell Physiology 36: 253–263.

Nievola CC, Carvalho CP, Carvalho V, Rodrigues E. 2017. Rapid responses of plants to temperature changes. Temperature 4: 371–405.

Obryk MK, Doran PT, Priscu JC. 2019. Prediction of Ice-Free Conditions for a Perennially Ice-Covered Antarctic Lake. Journal of Geophysical Research: Earth Surface 124: 686–694.

Pascual LS, Mohanty D, Sinha R, Nguyen TT, Rowland L, Lyu Z, Joshi T, Mooney BP, Gómez-Cadenas A, Fritschi FB, et al. 2025. Enhanced cell aggregation in the Chlamydomonas reinhardtii rbo1 mutant in response to multifactorial stress combination. Plant Physiology 199: kiaf551.

Patriarche JD, Priscu JC, Takacs-Vesbach C, Winslow L, Myers KF, Buelow H, Morgan-Kiss RM, Doran PT. 2021. Year-Round and Long-Term Phytoplankton Dynamics in Lake Bonney, a Permanently Ice-Covered Antarctic Lake. Journal of Geophysical Research: Biogeosciences 126: e2020JG005925.

Pocock T, Lachance M-A, Pröschold T, Priscu JC, Kim SS, Huner NPA. 2004. IDENTIFICATION OF A PSYCHROPHILIC GREEN ALGA FROM LAKE BONNEY ANTARCTICA: *CHLAMYDOMONAS RAUDENSIS* ETTL. (UWO 241) *CHLOROPHYCEAE*. Journal of Phycology 40: 1138–1148.

Pocock T, Vetterli A, Falk S. 2011. Evidence for phenotypic plasticity in the Antarctic extremophile Chlamydomonas raudensis Ettl. UWO 241. Journal of Experimental Botany 62: 1169–1177.

Poirier MC, Fugard K, Cvetkovska M. 2025. Light quality affects chlorophyll biosynthesis and photosynthetic performance in Antarctic Chlamydomonas. Photosynthesis Research 163: 9.

Popson D, D’Silva S, Wheeless K, Morgan-Kiss R. 2024. Permanent Stress Adaptation and Unexpected High Light Tolerance in the Shade-Adapted Chlamydomonas priscui. Plants 13: 2254.

Possmayer M, Berardi G, Beall BFN, Trick CG, Hüner NPA, Maxwell DP. 2011. PLASTICITY OF THE PSYCHROPHILIC GREEN ALGA CHLAMYDOMONAS RAUDENSIS (UWO 241) (CHLOROPHYTA) TO SUPRAOPTIMAL TEMPERATURE STRESS1: HEAT STRESS IN A PSYCHROPHILIC GREEN ALGA. Journal of Phycology 47: 1098–1109.

Possmayer M, Gupta RK, Szyszka-Mroz B, Maxwell DP, Lachance M, Hüner NPA, Smith DR. 2016. Resolving the phylogenetic relationship between *Chlamydomonas* sp. UWO 241 and *Chlamydomonas raudensis* sag 49.72 (Chlorophyceae) with nuclear and plastid DNA sequences (M Schroda, Ed.). Journal of Phycology 52: 305–310.

Pushpavalli R, Berger JD, Turner NC, Siddique KHM, Colmer TD, Vadez V. 2020. Cross-tolerance for drought, heat and salinity stresses in chickpea (Cicer arietinum L.). Journal of Agronomy and Crop Science 206: 405–419.

Rappaport HB, Oliverio AM. 2023. Extreme environments offer an unprecedented opportunity to understand microbial eukaryotic ecology, evolution, and genome biology. Nature Communications 14: 4959.

Ratcliff WC, Herron MD, Howell K, Pentz JT, Rosenzweig F, Travisano M. 2013. Experimental evolution of an alternating uni- and multicellular life cycle in Chlamydomonas reinhardtii. Nature Communications 4: 2742.

Rheinlænder JW, Regan H, Rampal P, Boutin G, Ólason E, Davy R. 2024. Breaking the Ice: Exploring the Changing Dynamics of Winter Breakup Events in the Beaufort Sea. Journal of Geophysical Research: Oceans 129: e2023JC020395.

Rillig MC, Ryo M, Lehmann A, Aguilar-Trigueros CA, Buchert S, Wulf A, Iwasaki A, Roy J, Yang G. 2019. The role of multiple global change factors in driving soil functions and microbial biodiversity. Science 366: 886–890.

Rivero RM, Mestre TC, Mittler R, Rubio F, Garcia-Sanchez F, Martinez V. 2014. The combined effect of salinity and heat reveals a specific physiological, biochemical and molecular response in tomato plants. Plant, Cell & Environment 37: 1059–1073.

Roland TP, Bartlett OT, Charman DJ, Anderson K, Hodgson DA, Amesbury MJ, Maclean I, Fretwell PT, Fleming A. 2024. Sustained greening of the Antarctic Peninsula observed from satellites. Nature Geoscience.

Saha M, Stroeve J, Isleifson D, Yackel J, Nandan V, Landy JC, Lam HM. 2025. Snow depth estimation on leadless landfast ice using Cryo2Ice satellite observations. The Cryosphere 19: 325–346.

Schroda M, Hemme D, Mühlhaus T. 2015. The *Chlamydomonas* heat stress response. The Plant Journal 82: 466–480.

Segawa T, Matsuzaki R, Takeuchi N, Akiyoshi A, Navarro F, Sugiyama S, Yonezawa T, Mori H. 2018. Bipolar dispersal of red-snow algae. Nature Communications 9: 3094.

Segawa T, Yonezawa T, Matsuzaki R, Mori H, Akiyoshi A, Navarro F, Fujita K, Aizen VB, Li Z, Mano S, et al. 2023. Evolution of snow algae, from cosmopolitans to endemics, revealed by DNA analysis of ancient ice. The ISME Journal 17: 491–501.

Shaar-Moshe L, Blumwald E, Peleg Z. 2017. Unique Physiological and Transcriptional Shifts under Combinations of Salinity, Drought, and Heat. Plant Physiology 174: 421–434.

Sherwell S, Kalra I, Li W, McKnight DM, Priscu JC, Morgan-Kiss RM. 2022. Antarctic lake phytoplankton and bacteria from near-surface waters exhibit high sensitivity to climate-driven disturbance. Environmental Microbiology 24: 6017–6032.

Silvano A, Rintoul SR, Peña-Molino B, Hobbs WR, van Wijk E, Aoki S, Tamura T, Williams GD. 2018. Freshening by glacial meltwater enhances melting of ice shelves and reduces formation of Antarctic Bottom Water. Science Advances 4: eaap9467.

Slater T, Lawrence IR, Otosaka IN, Shepherd A, Gourmelen N, Jakob L, Tepes P, Gilbert L, Nienow P. 2021. Review article: Earth’s ice imbalance. The Cryosphere 15: 233–246.

Smoot CA, Hopcroft RR. 2017. Depth-stratified community structure of Beaufort Sea slope zooplankton and its relations to water masses. Journal of Plankton Research 39: 79–91.

Soto DF, Gómez I, Huovinen P. 2023. Antarctic snow algae: unraveling the processes underlying microbial community assembly during blooms formation. Microbiome 11: 200.

Spolaor A, Scoto F, Larose C, Barbaro E, Burgay F, Bjorkman MP, Cappelletti D, Dallo F, de Blasi F, Divine D, et al. 2024. Climate change is rapidly deteriorating the climatic signal in Svalbard glaciers. The Cryosphere 18: 307–320.

Stahl-Rommel S, Kalra I, D’Silva S, Hahn MM, Popson D, Cvetkovska M, Morgan-Kiss RM. 2021. Cyclic electron flow (CEF) and ascorbate pathway activity provide constitutive photoprotection for the photopsychrophile, Chlamydomonas sp. UWO 241 (renamed Chlamydomonas priscuii). Photosynthesis Research.

Stibal M, Elster J, Sabacká M, Kastovská K. 2007. Seasonal and diel changes in photosynthetic activity of the snow alga Chlamydomonas nivalis (Chlorophyceae) from Svalbard determined by pulse amplitude modulation fluorometry. FEMS microbiology ecology 59: 265–273.

Stroeve J, Notz D. 2018. Changing state of Arctic sea ice across all seasons. Environmental Research Letters 13: 103001.

Sun Z, Cunningham FX, Gantt E. 1998. Differential expression of two isopentenyl pyrophosphate isomerases and enhanced carotenoid accumulation in a unicellular chlorophyte. Proceedings of the National Academy of Sciences of the United States of America 95: 11482–11488.

Suwannachuen N, Leetanasaksakul K, Roytrakul S, Phaonakrop N, Thaisakun S, Roongsattham P, Jantasuriyarat C, Sanevas N, Sirikhachornkit A. 2023. Palmelloid Formation and Cell Aggregation Are Essential Mechanisms for High Light Tolerance in a Natural Strain of Chlamydomonas reinhardtii. International Journal of Molecular Sciences 24: 8374.

Suzuki H, Hulatt CJ, Wijffels RH, Kiron V. 2019. Growth and LC-PUFA production of the cold-adapted microalga Koliella antarctica in photobioreactors. Journal of Applied Phycology 31: 981–997.

Szyszka B, Ivanov AG, Hüner NPA. 2007. Psychrophily is associated with differential energy partitioning, photosystem stoichiometry and polypeptide phosphorylation in Chlamydomonas raudensis. Biochimica et Biophysica Acta (BBA) - Bioenergetics 1767: 789–800.

Szyszka-Mroz B, Ivanov AG, Trick CG, Hüner NPA. 2022. Palmelloid formation in the Antarctic psychrophile, Chlamydomonas priscuii, is photoprotective. Frontiers in Plant Science 13: 911035.

Takizawa K, Takahashi S, Hüner NPA, Minagawa J. 2009. Salinity affects the photoacclimation of Chlamydomonas raudensis Ettl UWO241. Photosynthesis Research 99: 195–203.

Taylor RL, Semeniuk DM, Payne CD, Zhou J, Tremblay J-É, Cullen JT, Maldonado MT. 2013. Colimitation by light, nitrate, and iron in the Beaufort Sea in late summer. Journal of Geophysical Research: Oceans 118: 3260–3277.

Tonboe RT, Nandan V, Yackel J, Kern S, Pedersen LT, Stroeve J. 2021. Simulated Ka- and Ku-band radar altimeter height and freeboard estimation on snow-covered Arctic sea ice. The Cryosphere 15: 1811–1822.

Tremblay J-É, Robert D, Varela DE, Lovejoy C, Darnis G, Nelson RJ, Sastri AR. 2012. Current state and trends in Canadian Arctic marine ecosystems: I. Primary production. Climatic Change 115: 161–178.

Tremblay J-É, Simpson K, Martin J, Miller L, Gratton Y, Barber D, Price NM. 2008. Vertical stability and the annual dynamics of nutrients and chlorophyll fluorescence in the coastal, southeast Beaufort Sea. Journal of Geophysical Research: Oceans 113.

Turner J, Lu H, King J, Marshall GJ, Phillips T, Bannister D, Colwell S. 2021. Extreme Temperatures in the Antarctic.

Urbański JA, Litwicka D. 2022. The decline of Svalbard land-fast sea ice extent as a result of climate change. Oceanologia 64: 535–545.

Vyverman W, Verleyen E, Sabbe K, Vanhoutte K, Sterken M, Hodgson DA, Mann DG, Juggins S, Van de Vijver B, Jones V, et al. 2007. Historical processes constrain patterns in global diatom diversity. Ecology 88: 1924–1931.

Williamson CJ, Cook J, Tedstone A, Yallop M, McCutcheon J, Poniecka E, Campbell D, Irvine-Fynn T, McQuaid J, Tranter M, et al. 2020. Algal photophysiology drives darkening and melt of the Greenland Ice Sheet. Proceedings of the National Academy of Sciences 117: 5694–5705.

Wolf KKE, Hoppe CJM, Rehder L, Schaum E, John U, Rost B. 2024. Heatwave responses of Arctic phytoplankton communities are driven by combined impacts of warming and cooling. Science Advances 10: eadl5904.

Wunderling N, Willeit M, Donges JF, Winkelmann R. 2020. Global warming due to loss of large ice masses and Arctic summer sea ice. Nature Communications 11: 5177.

Young JN, Schmidt K. 2020. It’s what’s inside that matters: physiological adaptations of high-latitude marine microalgae to environmental change. New Phytologist 227: 1307–1318.

Zahra N, Hafeez MB, Ghaffar A, Kausar A, Zeidi MA, Siddique KHM, Farooq M. 2023. Plant photosynthesis under heat stress: Effects and management. Environmental and Experimental Botany 206: 105178.

Zandalinas SI, Mittler R. 2022. Plant responses to multifactorial stress combination. New Phytologist 234: 1161–1167.

Zandalinas SI, Sengupta S, Fritschi FB, Azad RK, Nechushtai R, Mittler R. 2021. The impact of multifactorial stress combination on plant growth and survival. New Phytologist 230: 1034–1048.

Zhang X, Cvetkovska M, Morgan-Kiss R, Hüner NPA, Smith DR. 2021. Draft genome sequence of the Antarctic green alga Chlamydomonas sp. UWO241. *iScience* 24: 102084.

Zhang Z, Qu C, Yao R, Nie Y, Xu C, Miao J, Zhong B. 2019. The Parallel Molecular Adaptations to the Antarctic Cold Environment in Two Psychrophilic Green Algae. Genome Biology and Evolution 11: 1897–1908.

Zhang Z, Qu C, Zhang K, He Y, Zhao X, Yang L, Zheng Z, Ma X, Wang X, Wang W, et al. 2020. Adaptation to Extreme Antarctic Environments Revealed by the Genome of a Sea Ice Green Alga. Current Biology 30: 3330–3341.e7.

