## Supporting information for "Temperature stress resilience in polar *Chlamydomonas* is regulated by acclimation to light and salinity: implications for survival in a changing world"

### Supplementary Information

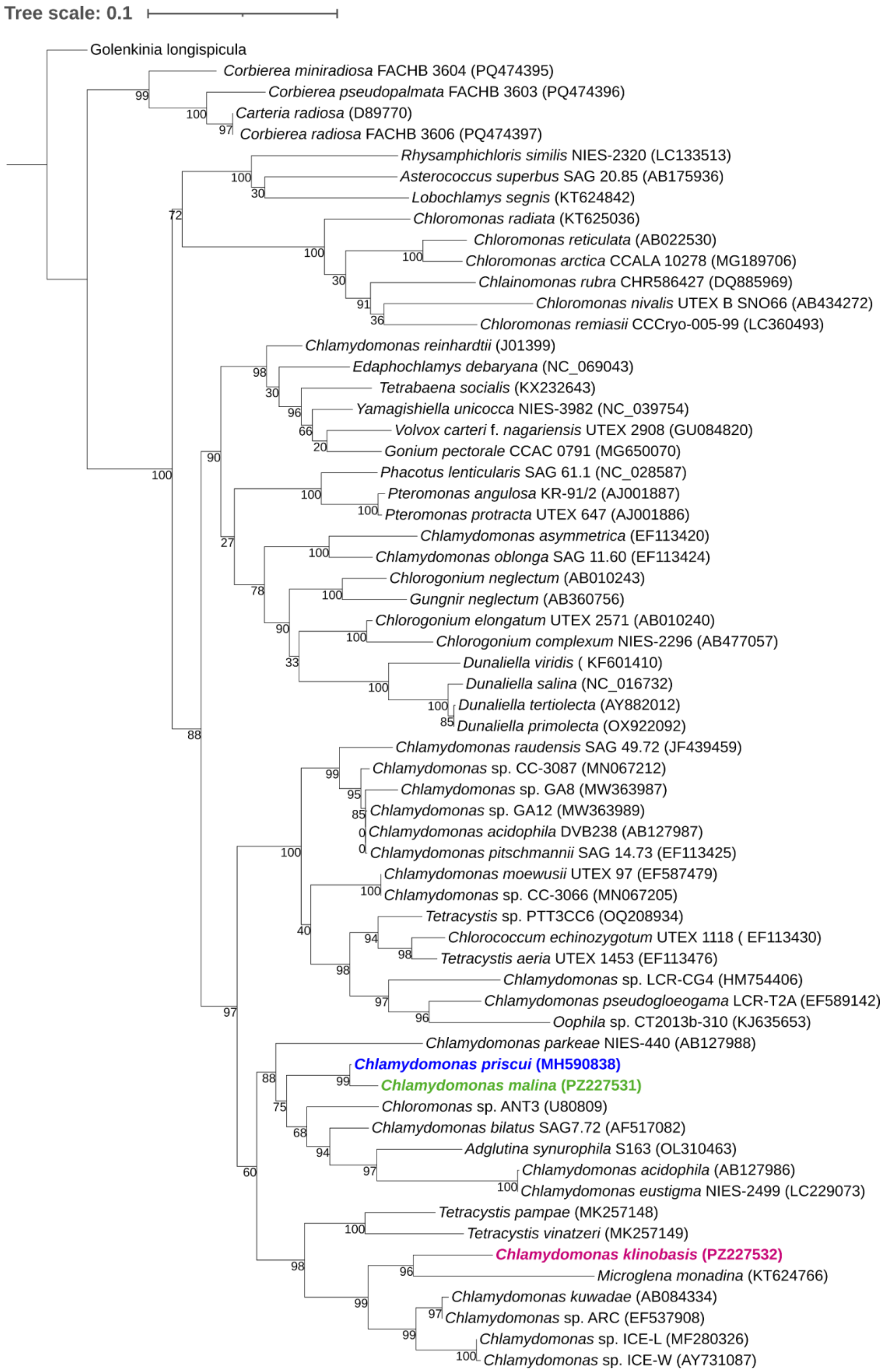


**Figure S1:** Phylogenetic tree of 63 rbcL nucleotide sequences inferred from using maximum likelihood analysis is shown to scale (substitutions per site), with branch values representing bootstrap support. GenBank accession numbers are provided in brackets beside the organism name. The species considered in this study are highlighted in blue (*Chlamydomonas priscui*), green (*Chlamydomonas malina*) and mauve (*Chlamydomonas klinobasis*).

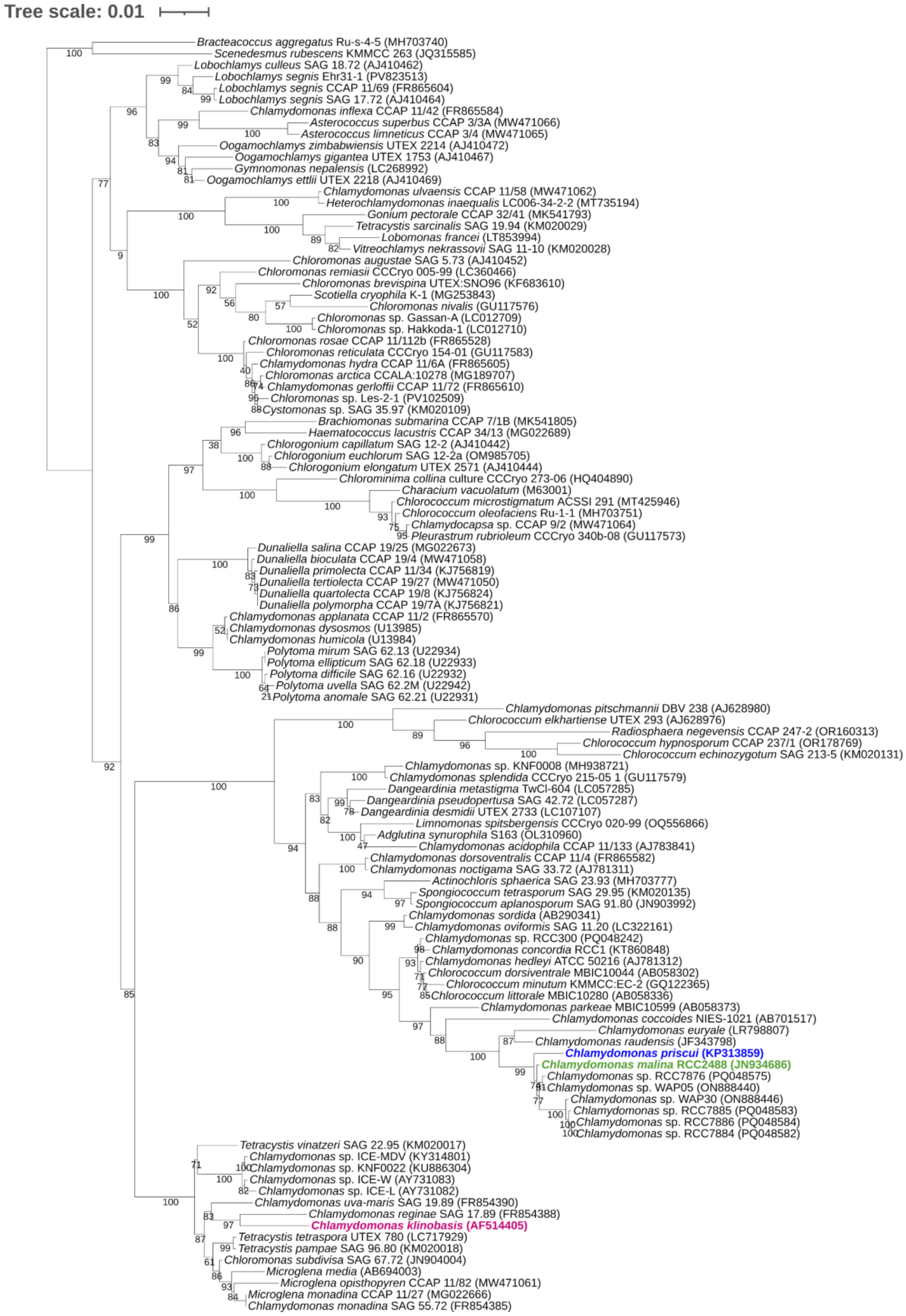


**Figure S2:** Phylogenetic tree of 111 18S rRNA sequences inferred from using maximum likelihood analysis is shown to scale (substitutions per site), with branch values representing bootstrap support. GenBank accession numbers are provided in brackets beside the organism name. The species considered in this study are highlighted in blue (*Chlamydomonas priscui*), green (*Chlamydomonas malina*) and mauve (*Chlamydomonas klinobasis*).

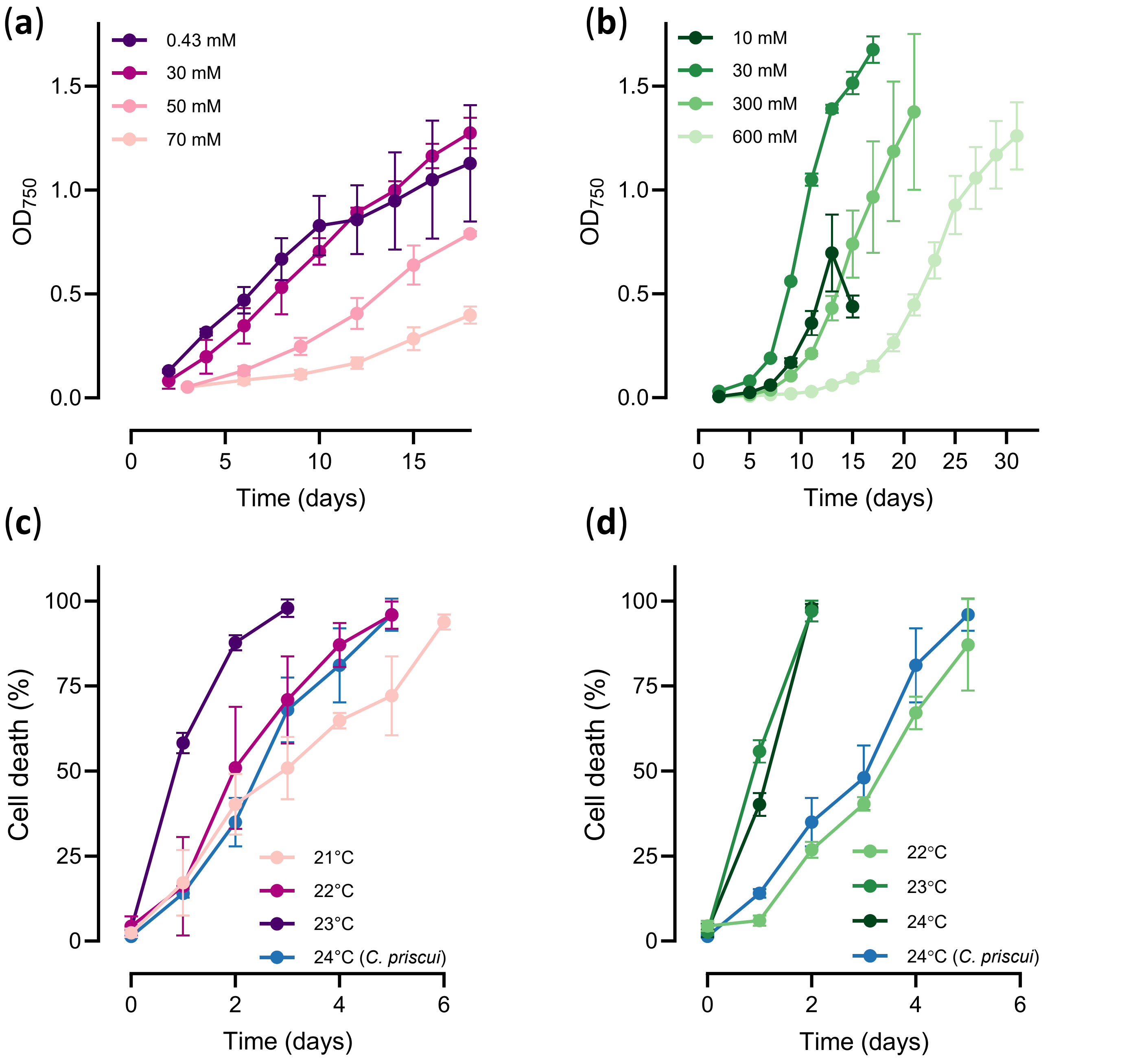


**Figure S3**: Optimizing the growth and stress conditions in *C. klinobasis* and *C. malina*. (**a**) Optical density measured at a range of salinities (0.43-70 mM NaCl) in *C. klinobasis* cultures. (**b**) Optical densities measured at a range of salinities (10-600 mM) in *C. malina* cultures. In both (a) and (b), algal cultures cultured at 4°C. (c) Kinetics of cell death in *C. klinobasis* exposed to non-permissive temperatures (21°C, 22°C, 23°C). (d) Kinetics of cell death in *C. malina* exposed to non-permissive temperatures (22°C, 23°C, 24°C). In both (c) and (d), the algae were acclimated to the lowest salinity that supports robust growth (0.43 mM and 30 mM NaCl for *C. klinobasis* and *C. malina*, respectively). The blue line corresponds to the kinetics of cell death in *C. priscui* acclimated to 10 mM NaCl and exposed to 24°C. In all cases, algae were cultured at a constant light intensity of 100 µmol $m^{-2}$ $s^{-1}$. Data are means ± SD of at least three independent experiments.

**Table S1**: A comparison of the growth rates in *C. priscui* (P), *C. malina* (M), and *C. klinobasis* (K) cultured at 4°C and acclimated to high salinity (HS), low salinity (LS), high light (HL) or low light (LL). Data are means ± SD of at least three biological replicates. The lack of statistical difference ($p > 0.05$) between the growth rates of the three species was confirmed with a one-way ANOVA with a Tukey's post-hoc test preformed within each treatment.

| | **Growth rate ($\Delta OD_{750}$ $d^{-1}$)** | | | ***p*-value** | | |
|---|---|---|---|---|---|---|
| **Treatment** | ***C. priscui* (P)** | ***C. malina* (M)** | ***C. klinobasis* (K)** | **P:M** | **P:K** | **M:K** |
| **HS-LL** | 0.19±0.03 | 0.21±0.02 | 0.15±0.02 | 0.804 | 0.454 | 0.0931 |
| **HS-HL** | 0.35±0.14 | 0.45±0.07 | 0.28±0.03 | 0.3779 | 0.5934 | 0.1291 |
| **LS-LL** | 0.26±0.05 | 0.23±0.03 | 0.23±0.02 | 0.7477 | 0.6947 | 0.9947 |
| **LS-HL** | 0.49±0.03 | 0.71±0.08 | 0.47±0.03 | 0.121 | 0.7102 | 0.0919 |

**Table S2**: A statistical summary of two-way ANOVA analyses for the interaction effects between the experimental variables (salinity, light intensity) and the measured morphological and physiological parameters in *C. priscui*, *C. malina* and *C. klinobasis.* The numbers represent p-values, and significant interactions ($p < 0.05$) are bolded.

| | **Parameter** | | | | | |
|---|---|---|---|---|---|---|
| **Variable** | **Growth rate** | **Cell size (μm)** | **Palmelloids (%)** | **Chl/cell (pg)** | **Chl a/b** | **Car/Chl** |
| | | | ***C. priscui*** | | | |
| Salinity | 0.1319 | 0.2599 | 0.1950 | 0.6016 | 0.0668 | **0.0113** |
| Light Intensity | **0.0063** | **0.0031** | **<0.001** | **<0.001** | **<0.001** | **<0.001** |
| Salinity:Light Intensity | 0.5114 | 0.3384 | 0.1365 | 0.8932 | 0.7794 | 0.7785 |
| | | | ***C. malina*** | | | |
| Salinity | **0.0031** | 0.1272 | 0.2094 | 0.5759 | **<0.001** | 0.0029 |
| Light Intensity | **<0.001** | **0.0226** | **<0.001** | **0.0248** | **<0.001** | **<0.001** |
| Salinity:Light Intensity | **0.0093** | 0.6578 | 0.3184 | 0.3833 | 0.1458 | 0.5104 |
| | | | ***C. klinobasis*** | | | |
| Salinity | **<0.001** | 0.0908 | 0.7317 | 0.0469 | 0.1983 | **<0.001** |
| Light Intensity | **<0.001** | **<0.001** | **<0.001** | **0.0216** | **<0.001** | **<0.001** |
| Salinity:Light Intensity | **0.0049** | 0.4582 | 0.0278 | 0.1443 | 0.3813 | **0.0067** |

Chl, chlorophyll; Car, carotenoids

**Table S3**: A statistical summary of three-way ANOVA analyses for the interaction effects between the experimental variables (salinity, light intensity, time at non-permissive temperature) and the measured viability and physiological parameters in *C. priscui*, *C. malina* and *C. klinobasis.* The numbers represent p-values, and significant interactions ($p < 0.05$) are bolded.

| | Parameter | | | | | | |
|---|---|---|---|---|---|---|---|
| **Variable** | **Cell death (%)** | **Chl loss (%)** | **$F_V/F_M$** | **ETR (µmol $m^{-2}$ $s^{-1}$)** | **Y(II)** | **Y(NPQ)** | **Y(NO)** |
| ***C. priscui*** | | | | | | | |
| Salinity | 0.1220 | 0.4420 | **0.0410** | 0.0860 | 0.0860 | **0.0330** | 0.0690 |
| Light Intensity | 0.9520 | 0.3670 | **0.0280** | 0.9250 | 0.9250 | 0.9100 | 0.3920 |
| Time | **<0.001** | **<0.001** | **<0.001** | **<0.001** | **<0.001** | **<0.001** | **<0.001** |
| Salinity:Light Intensity | 0.8000 | 0.8710 | 0.5300 | 0.0700 | 0.0710 | 0.1740 | 0.7430 |
| Salinity:Time | **<0.001** | **<0.001** | **<0.001** | **<0.001** | **<0.001** | **<0.001** | 0.1160 |
| Light Intensity:Time | 0.0874 | **0.0102** | **0.0010** | **0.0030** | **0.0030** | **0.0230** | 0.0720 |
| Salinity:Light Intensity:Time | **<0.001** | **<0.001** | 0.1410 | **0.0250** | **0.0240** | 0.0920 | 0.5620 |
| ***C. malina*** | | | | | | | |
| Salinity | 0.2650 | 0.1650 | 0.0590 | **0.0410** | **0.0410** | 0.1470 | 0.0900 |
| Light Intensity | 0.5220 | 0.7940 | 0.0510 | 0.1640 | 0.1640 | 0.4170 | 0.3400 |
| Time | **<0.001** | **<0.001** | **0.0005** | **<0.001** | **<0.001** | **0.0010** | **0.0003** |
| Salinity:Light Intensity | 0.9400 | 0.2370 | 0.1890 | 0.0540 | 0.0530 | 0.8900 | 0.0760 |
| Salinity:Time | **<0.001** | **0.0023** | **0.0002** | **0.0020** | **0.0020** | **0.0050** | **0.0260** |
| Light Intensity:Time | **<0.001** | **<0.001** | **0.0010** | **0.0130** | **0.0120** | 0.0670 | 0.0620 |
| Salinity:Light Intensity:Time | 0.3780 | 0.1250 | **0.0100** | **0.0180** | **0.0180** | 0.2030 | 0.2310 |
| ***C. klinobasis*** | | | | | | | |
| Salinity | 0.9260 | 0.1993 | **0.0410** | 0.0860 | 0.0860 | **0.0330** | 0.0690 |
| Light Intensity | 0.6050 | **0.0032** | **0.0280** | 0.9250 | 0.9250 | 0.9100 | 0.3920 |
| Time | **<0.001** | **<0.001** | **<0.001** | **<0.001** | **<0.001** | **<0.001** | **<0.001** |
| Salinity:Light Intensity | 0.1820 | 0.8577 | 0.5300 | 0.0700 | 0.0710 | 0.1740 | 0.7430 |
| Salinity:Time | **<0.001** | 0.1037 | **<0.001** | **<0.001** | **<0.001** | **<0.001** | 0.1160 |
| Light Intensity:Time | **<0.001** | **<0.001** | **0.0010** | **0.0030** | **0.0030** | **0.0230** | 0.0720 |
| Salinity:Light Intensity:Time | **0.0088** | 0.6150 | 0.1410 | **0.0250** | **0.0240** | 0.0920 | 0.5620 |

Chl, chlorophyll; $F_V/F_M$, maximum PSII photochemical efficiency; ETR, relative electron transport rate at PSII; Y(II), photochemical yield of PSII; Y(NPQ) non-photochemical quenching; Y(NO) non-regulated dissipation

| | | Energy Partitioning | | |
|---|---|---|---|---|
| **Treatment** | **Time (h)** | **Y(II)** | **Y(NO)** | **Y(NPQ)** |
| | | ***C. priscui*** | | |
| **HS-LL** | **0** | 0.313±0.073 | 0.556±0.072 | 0.13±0.002 |
| | **6** | 0.388±0.047 (ns) | 0.481±0.048 (ns) | 0.131±0.001 (ns) |
| | **24** | 0.411±0.016 (ns) | **0.446±0.007 (*)** | 0.143±0.01 (ns) |
| | **48** | **0.347±0.02 (**)** | 0.481±0.007 (ns) | **0.172±0.017 (**)** |
| **HS-HL** | **0** | 0.351±0.039 | 0.534±0.041 | 0.115±0.002 |
| | **6** | 0.451±0.028 (ns) | **0.445±0.018 (*)** | 0.105±0.01 (ns) |
| | **24** | **0.328±0.016 (*)** | 0.511±0.02 (ns) | **0.161±0.011 (*)** |
| | **48** | **0.24±0.003 (***)** | 0.542±0.028 (ns) | **0.217±0.029 (***)** |
| **LS-LL** | **0** | 0.518±0.035 | 0.354±0.015 | 0.128±0.025 |
| | **6** | 0.468±0.027 (ns) | 0.384±0.012 (ns) | 0.148±0.017 (ns) |
| | **24** | 0.319±0.073 (ns) | **0.457±0.013 (***)** | 0.225±0.084 (ns) |
| | **48** | **0.117±0.043 (***)** | **0.537±0.033 (****)** | **0.346±0.018 (***)** |
| **LS-HL** | **0** | 0.555±0.036 | 0.337±0.026 | 0.108±0.016 |
| | **6** | 0.575±0.032 (ns) | 0.315±0.018 (ns) | 0.111±0.014 (ns) |
| | **24** | **0.36±0.041 (**)** | **0.422±0.024 (**)** | **0.218±0.054 (**)** |
| | **48** | **0±0 (****)** | **0.612±0.025 (****)** | **0.388±0.025 (****)** |
| | | ***C. malina*** | | |
| **HS-LL** | **0** | 0.481±0.014 | 0.425±0.009 | 0.094±0.006 |
| | **6** | 0.445±0.024 (ns) | 0.419±0.017 (ns) | **0.137±0.008 (****)** |
| | **24** | **0.386±0.009 (***)** | **0.462±0.006 (*)** | **0.152±0.003 (****)** |
| | **48** | **0.27±0.005 (****)** | **0.524±0.006 (****)** | **0.206±0.001 (****)** |
| **HS-HL** | **0** | 0.528±0.043 | 0.412±0.052 | 0.06±0.011 |
| | **6** | **0.417±0.032 (*)** | 0.444±0.032 (ns) | **0.14±0.002 (*)** |
| | **24** | **0.227±0.047 (****)** | **0.544±0.025 (**)** | **0.229±0.041 (***)** |
| | **48** | **0.056±0.022 (****)** | **0.547±0.012 (*)** | **0.398±0.034 (****)** |
| **LS-LL** | **0** | 0.472±0.066 | 0.416±0.052 | 0.113±0.014 |
| | **6** | **0.342±0.02 (*)** | 0.457±0.021 (ns) | 0.201±0.015 (ns) |
| | **24** | **0.154±0.036 (****)** | 0.46±0.045 (ns) | **0.386±0.078 (***)** |
| | **48** | **0±0 (****)** | **0.711±0.027 (****)** | **0.289±0.027 (**)** |
| **LS-HL** | **0** | 0.579±0.022 | 0.377±0.018 | 0.045±0.004 |
| | **6** | **0.366±0.017 (****)** | **0.507±0.05 (*)** | 0.127±0.033 (ns) |
| | **24** | **0±0.001 (****)** | **0.649±0.051 (***)** | **0.35±0.051 (****)** |
| | **48** | **0±0 (****)** | **0.698±0.032 (****)** | **0.302±0.032 (***)** |
| | | ***C. klinobasis*** | | |
| **HS-LL** | **0** | 0.5±0.007 | 0.351±0.011 | 0.149±0.004 |
| | **6** | **0.555±0.015 (*)** | **0.307±0.009 (*)** | 0.138±0.006 (ns) |
| | **24** | 0.539±0.004 (ns) | 0.331±0.003 (ns) | **0.131±0.005 (*)** |
| | **48** | 0.49±0.037 (ns) | 0.377±0.026 (ns) | 0.133±0.011 (ns) |
| **HS-HL** | **0** | 0.43±0.039 | 0.392±0.018 | 0.178±0.021 |
| | **6** | **0.523±0.029 (**)** | **0.346±0.02 (**)** | **0.131±0.009 (*)** |
| | **24** | **0.504±0.005 (*)** | **0.358±0.003 (*)** | **0.138±0.005 (*)** |
| | **48** | 0.428±0.016 (ns) | 0.391±0.006 (ns) | 0.181±0.021 (ns) |
| **LS-LL** | **0** | 0.483±0.017 | 0.385±0.025 | 0.133±0.008 |
| | **6** | 0.531±0.022 (ns) | 0.342±0.019 (ns) | 0.128±0.006 (ns) |
| | **24** | 0.483±0.018 (ns) | 0.383±0.015 (ns) | 0.135±0.003 (ns) |
| | **48** | **0.404±0.026 (**)** | **0.449±0.023 (*)** | **0.147±0.005 (*)** |
| **LS-HL** | **0** | 0.513±0.006 | 0.382±0.003 | 0.105±0.007 |
| | **6** | **0.569±0.03 (*)** | **0.314±0.032 (*)** | 0.117±0.004 (ns) |
| | **24** | 0.478±0.021 (ns) | 0.364±0.025 (ns) | **0.158±0.006 (**)** |
| | **48** | **0.318±0.027 (****)** | **0.448±0.008 (*)** | **0.234±0.02 (****)** |

**Table S4**: A statistical summary of three-way ANOVA analyses followed by Dunnett's post hoc test for the energy partitioning parameters measured in *C. prisui*, *C. malina* and *C. klinobasis*. The total absorbed light is distributed between photochemical yield of PSII Y(II), non-photochemical quenching Y(NPQ), and non-regulated dissipation Y(NO). In all cases, algae were cultured at 4°C and acclimated to high salinity (HS), low salinity (LS), high light (HL) or low light (LL). Temperature stress was initiated at the mid-exponential growth phase (OD750 = 0.3-0.4) and corresponds to t = 0 hours. Data are means ± SD of at least three biological replicates. Statistical significance is indicated by an asterisk and bolded (* = < 0.05, ** = < 0.01, *** = < 0.001, **** = < 0.0001).
